# *Enterobacter* sp. SA187-induced coordinated regulation of high-affinity nitrate transporters and ethylene signaling enhances nitrogen content and plant growth under low nitrate

**DOI:** 10.1101/2025.06.23.660384

**Authors:** Amina Ilyas, Caroline Mauve, Berengere Decouard, Jose Caius, Christine Paysant-Leroux, Michael Hodges, Axel de Zelicourt

## Abstract

- Sustainable crop production demands solutions to reduce the overuse of synthetic nitrogen fertilizers, and plant-growth-promoting bacteria offer a promising strategy by enhancing nutrients acquisition. This study investigated ability of a non-diazotrophic bacterium, *Enterobacter* sp. SA187 (SA187), in enhancing Arabidopsis growth under low nitrate conditions and the underlying mechanisms.
- Arabidopsis seedlings were grown under different nitrate concentrations with or without SA187 inoculation. Growth traits were quantified alongside shoot and root nitrate and total nitrogen contents, and C:N ratios. Transcriptomic profiling (RNA-seq) and qRT-PCR were used to assess modified gene expression. Functional validation was conducted using ethylene-insensitive (*ein2-1*) and high-affinity nitrate transporter (HATS) mutants (*nrt2.5, nrt2.6*).
- SA187 significantly enhanced fresh weight, primary root length, and lateral root density under low nitrate, with benefits increasing as nitrate availability decreased. SA187 improved nitrate accumulation and shoot nitrogen allocation, reducing shoot C:N ratios. SA187 regulated expression of HATS and hormone-responsive genes. The growth-promoting effects were abolished in *ein2-1, nrt2.5,* and *nrt2.6* mutants, and SA187-induced regulation of *NRT2.5* occurred downstream of ethylene signaling, while *NRT2.6* was partly ethylene-independent.
- SA187 promotes growth under low nitrate possibly through ethylene-mediated and HATS-dependent reprogramming of nitrate accumulation and nitrogen allocation, supporting its use as a microbial solution for low-input agriculture.

## I. Introduction

Nitrogen (N) is essential for plant growth, development, and productivity. As a core component of nucleic acids, proteins, amino acids, chlorophyll, and secondary metabolites, N coordinates numerous physiological and metabolic processes (Krapp, 2015; O’Brien *et al*., 2016). Consequently, N-deficiency results in chlorosis, reduced photosynthetic capacity, stunted growth, and yield losses (Ding *et al*., 2005; Hawkesford *et al*., 2012; Ahmad Wani *et al*., 2022). Although atmospheric nitrogen (N_2_) is abundant, most plants cannot directly use this inert form, instead they rely on bioavailable N-compounds such as nitrate (NO_3_^-^), ammonium (NH_4_^+^), and amino acids. Among these, NO_3_^-^ generally predominates as the available N-form in well-aerated soils (Glass, 2009; Krapp *et al*., 2014). However, the availability of bioavailable N-compounds is often limited in agricultural systems thus making this a significant factor limiting crop yields worldwide (LeBauer & Treseder, 2008; Sinclair & Rufty, 2012).

To alleviate N-limitations and ensure global food security, agriculture has relied on synthetic N-based fertilizers. While this has significantly improved crop yields, their use remains inefficient, costly and they are a source of greenhouse gases and pollution. N-use-efficiency (NUE) remains remarkably low as less than 50% of applied N is taken up by plants, while the remainder is lost to volatilization, leaching, and denitrification. These losses not only reduce productivity and economic returns for farmers but also cause widespread environmental degradation (Liu *et al*., 2010; Ali *et al*., 2025). Adverse impacts include groundwater nitrate contamination, eutrophication, greenhouse gas emissions (such as N_2_O), ozone depletion, and long-term soil degradation, with serious consequences for ecosystems and human health (FAO & UNEP, 2021; Sutton *et al*., 2021; Nath *et al*., 2023; Govindasamy *et al*., 2023; de Vries *et al*., 2024). In parallel, the production of N-based fertilizers via the Haber–Bosch process is energy-intensive and carbon (C)-emitting, adding further environmental and economic burdens (Cherkasov *et al*., 2015; Rouwenhorst & Lefferts, 2020). Therefore, there is a need to improve plant N-uptake and N-utilization so as to support plant growth under low-N-input conditions, reducing fertilizer use in the context of an environmentally-friendly, sustainable agriculture.

Plant-growth-promoting bacteria (PGPB) have emerged as promising biological tools to reduce reliance on synthetic inputs. Historically, research has extensively focused on symbiotic N_2_-fixing bacteria (e.g., *Rhizobia*, *Frankia* spp.). However, emerging studies demonstrate that even non-N_2_-fixing endophytic bacteria can significantly improve plant growth directly, by enhancing nutrient availability, mobilizing soil minerals such as phosphorus, potassium, iron and N, and indirectly, by modulating hormonal signaling pathways, root architecture, defense and systemic stress responses (Bakhshandeh et al., 2020; de Zelicourt et al., 2013; Di Benedetto et al., 2017; Etesami & Adl, 2020; Glick, 2012, 2020; Pathania et al., 2020; Ramakrishna et al., 2019). Despite increasing interest, the mechanistic details underlying these beneficial plant–microbe interactions, particularly under N-limiting conditions, remain insufficiently characterized.

Among promising non-diazotrophic plant-associated endophytes, *Enterobacter* sp. SA187 (SA187) has emerged as a particularly effective beneficial endophytic bacterium. Initially isolated from the desert-adapted legume *Indigofera argentea* (Andrés-Barrao *et al*., 2017), SA187 effectively enhances resilience across a wide spectrum of abiotic stresses, including salinity, heat, and high light. The beneficial effects of SA187 are mediated primarily through modulation of critical physiological and molecular processes such as hormonal homeostasis, particularly involving ethylene signaling pathways, maintenance of nutrient and redox balances, and activation of stress-responsive networks (de Zélicourt et al., 2018; Rahman et al., 2025; Rolli et al., 2022; Shekhawat et al., 2024; Shekhawat et al., 2021). Although previous studies have demonstrated plant growth improvement by *Enterobacter* species under nutrient-limited conditions, through mechanisms such as N_2_-fixation, improved nutrient acquisition and altered root system architecture (Shankar *et al*., 2011; Lin *et al*., 2012; Madhaiyan *et al*., 2013; Zhu *et al*., 2013; Tsegaye *et al*., 2022), the role of SA187 under low N conditions has not been investigated.

Nitrate uptake and allocation within higher plants are primarily regulated by specialized nitrate transporters belonging to two main families: the NRT1/PTR family (NPF) comprising mostly low-affinity transporters (LATs), and the NRT2 family of high-affinity nitrate transporters (HATS). NRT2 family members are important under low nitrate availability conditions since they transport nitrate at very low external concentrations (Krapp *et al*., 2014; Zhang *et al*., 2024). HATS regulate not only soil-derived nitrate uptake but also nitrate distribution within plant tissues and organs, significantly influencing NUE and plant adaptation under nitrate scarcity (Wang *et al*., 2012; Lezhneva *et al*., 2014; Fan *et al*., 2017). Moreover, hormonal signals, notably ethylene, have been implicated in modulating the expression and functional activity of nitrate transporters (Tian et al., 2009; Zhang et al., 2014; Zheng et al., 2013). Ethylene signaling, mediated through EIN2 and EIN3, plays a central role in integrating nutrient and environmental cues, but its role in regulating nitrate accumulation via plant–microbe interactions remains poorly understood (Iqbal et al., 2017; Khan et al., 2017; Leblanc et al., 2008). Although SA187 is known to modulate ethylene signaling to enhance stress tolerance (de Zélicourt *et al*., 2018; Shekhawat *et al*., 2024), to date its specific role on nitrate transporter regulation and plant N-metabolism has not been explored. Since ethylene is known to interact with several nitrate transporters (including NRT2.1) (Tian *et al*., 2009; Zheng *et al*., 2013), it is possible that SA187-mediated modulation of ethylene signaling could significantly influence nitrate uptake and nitrate accumulation under low nitrate conditions.

This study investigated the beneficial impact of SA187 on the growth of *Arabidopsis thaliana* seedlings with respect to nitrate concentration, with a focus on growth responses, nitrate transporter expression, nitrate content, C:N ratios and ethylene signaling. Using phenotypic, elemental, transcriptomic, and genetic analyses, including HATS and ethylene-insensitive mutants, this work revealed that SA187 appeared to improve nitrate uptake and aerial allocation through a combination of ethylene-dependent and -independent regulatory pathways. These findings provide mechanistic insights into how a beneficial endophyte can modulate plant nutrient physiology, offering a path forward for microbial strategies to potentially enhance NUE under N-limited conditions.

## II. Materials and Methods

### A. Bacterial and plant material

*Arabidopsis thaliana* (L.) Heynh., ecotype Columbia-0 (Col-0, wild-type), and selected mutants in a Col-0 background: *ein2-1* (Guzmán & Ecker, 1990), *ein3-1* (Chao *et al*., 1997), *nrt2.5* (GK 213H10), *nrt2.6* (SM 3.35179), and a *nrt2.5 x nrt2.6* double mutant (Kechid *et al*., 2013) were used in this study. The *Enterobacter* sp. SA187 strain, originally isolated from root nodules of *Indigofera argentea* in Jizan, Saudi Arabia (Andrés-Barrao *et al*., 2017), was used in its GFP-tagged form (de Zélicourt *et al*., 2018).

### B. Plant growth and bacterial colonization

*Arabidopsis thaliana* seeds were surface-sterilized and inoculated with SA187 as described previously (Saad *et al*., 2018). Briefly, sterilized seeds were sown on modified half-strength Murashige and Skoog (½MS) medium prepared from nitrogen-free MS basal salts (Apollo Scientific Catalogue No: PMM531B), supplemented with 0.5 g L ¹ MES (pH 5.8) and 9 g L ¹ agar. Nitrate regimes were established by adding 0.2, 1, 2, 10, or 20 mM KNO . To equalize potassium across treatments, KCl was added from a 2 M stock solution so that the total added K (from KNO + KCl) was adjusted to 20 mM. Per liter of medium, the required volumes of KCl stock were: 9.9 mL (0.2 mM KNO), 9.5 mL (1 mM), 9.0 mL (2 mM), 5.0 mL (10 mM), and none for 20 mM KNO . For SA187 treatments (+SA187), 2×10^5^ bacteria/mL were added directly to the growth media before pouring into square petri plates. For mock controls (Mock), an equal volume of sterile LB medium was added instead. After stratification for 24 h at 4°C in the dark, plates were placed vertically in a growth chamber (16-h photoperiod, 18–20°C night/day, 120 μE/m^2^/s light intensity, 50 ± 10% relative humidity) to allow seed germination.

At 6 days post-germination (dpg), seedlings were transferred to fresh modified ½MS agar plates containing the same nitrate concentrations as in the germination plates and harvested at 16 days for fresh weight (FW) measurements, nitrate quantification and, molecular and elemental analyses. Plates were scanned at 16dpg to obtain the images for primary root length (PRL) and lateral root density (LRD) measurements.

### C. Growth phenotyping

FW of individual plants was measured at harvest. PRL and LRD were measured from plate scans using Fiji/ImageJ software (Schindelin et al., 2012).

### D. Quantification of bacterial colonization

Bacterial colonization was quantified as described previously (Saad *et al*., 2018). Briefly, 16-day-old roots and shoots of *Arabidopsis thaliana* seedlings were harvested separately, weighed, and homogenized in sterile 10 mM MgCl . Serial dilutions of homogenates were plated on LB agar containing rifampicin (50 µg mL ¹) to selectively recover GFP-tagged SA187 colonies. Plates were incubated at 30 °C overnight, and colony-forming units (CFUs) were counted. Colonization levels were expressed as CFUs per mg fresh weight. Four independent biological replicates were performed for each genotype and organ.

### E. Elemental analyses

Dried root and shoot tissues (∼1 mg per sample) were transferred to tin capsules, weighed and combusted in a Pyrocube elemental analyzer (Elementar, Lyon, France). C and N contents were determined via thermal conductivity detection (TCD) after gas separation on selective columns. Quantification was performed against certified standards: ammonium sulfate (21.2% N), sucrose (42.11% C), glutamic acid (9.51% N, 40.82% C), and glutamine (19.17% N, 41.09% C). Data are presented as % dry weight (DW).

### F. Nitrate content assay

Flash-frozen tissues (≥10 mg FW) were ground under liquid N. Each sample was extracted with 500 µL preheated Milli-Q water (100°C), vortexed, and boiled for 15 min with intermittent mixing. After cooling and centrifugation (15,000×g, 15 min), 450 µL of the supernatant was collected. Nitrate content was measured colorimetrically using the Nitrite/Nitrate Assay Kit (Sigma-Aldrich, #23479), following the manufacturers protocol. Technical triplicates were performed per sample.

### G. Transcriptomics

#### i. RNA-sequencing and data analyses

Shoot and root samples were collected from 16-day-old mock and SA187-treated plants grown at 20 mM and 1 mM nitrate, with three biological replicates per condition. Each replicate comprised pooled shoot or root tissues from 4–6 plants. Total RNA was extracted using the NucleoSpin RNA kit following the manufacturer’s recommendations (Macherey-Nagel). RNA-seq libraries were prepared from 500 ng total RNA using the QuantSeq 3′ mRNA-Seq Library Prep Kit (Lexogen) and sequenced (75 bp single-end) on an Illumina NextSeq500.

Unique molecular identifiers (UMIs) were removed and appended to the read identifier with the extract command of UMI-tools (v1.0.1) (Smith et al., 2017). Reads where any UMI base quality score fell below 10 were removed. To remove adapter sequences, poly(A), poly(G) sequences, and low quality nucleotides, reads were trimmed with BBduk from the BBmap suite (v38.84) (Bushnell, 2014) with the options k=13 ktrim=r useshortkmers=t mink=5 qtrim=r trimq=10 minlength=30. Trimmed reads were then mapped onto the reference genome of *Arabidopsis thaliana* (TAIR10+ARAPORT11+MtCol0) and counted using STAR (v2.7.3a) (Dobin et al., 2013), with the following parameters --alignIntronMin 5 --alignIntronMax 60000 -- alignMatesGapMax 6000 –alignEndsType Local--outFilterMultimapNmax 20 -- outFilterMultimapScoreRange 0 –outSAMprimaryFlag AllBestScore –mismatchNoverLmax 0,6. Reads with identical mapping coordinates and UMI sequences were collapsed to remove PCR duplicates using the dedup command of UMI-tools with the default directional method parameter. Deduplicated reads were counted using HTSeq version v0.12.4 (Anders et al., 2015) (htseq-count mode intersection-nonempty) based on the *A. thaliana* TAIR10-ARAPORT11 annotation. Between 2.9 (86.6% of raw reads) and 2.5 (95% of raw reads) millions of reads were associated to annotated genes.

Statistical analyses were conducted with R v3.6.2 (R Core Team, 2020) using DiCoExpress V1 (Lambert *et al*., 2020; Baudry *et al*., 2022) based on the Bioconductor package edgeR (v 3.28.0) (Robinson *et al*., 2010; McCarthy *et al*., 2012). Two separate statistical analyses were conducted on data from roots and leaves, starting from the raw count tables.

For each analysis, genes with low counts were filtered using the “filterByExpr” function where the group argument specifies the biological conditions, the min.count value was set to 3 and the min.total.count, large.n and min.prop arguments were set to their default values. Libraries were normalized using the TMM method.

For both analyses, the differential analysis was based on a negative binomial generalized linear model in which the logarithm of the average gene expression is an additive function of a nitrate effect (1mM or 20mM), an inoculation effect (SA187 or mock), their interaction and a replicate effect (3 modalities). Four comparisons were performed (i) the difference between 1 mM and 20 mM nitrate in the mock condition, (ii) the difference between 1 mM and 20 mM nitrate in the SA187 inoculated condition, (iii) the difference between mock and SA187 inoculated at 1 mM nitrate, and (iv) the difference between mock and SA187 inoculated at 20 mM nitrate. For each contrast a likelihood ratio test was applied and raw p-values were adjusted with the Benjamini– Hochberg procedure to control the false discovery rate. The distribution of the resulting p-values followed the quality criterion described by Rigaill et al. (2016). A gene was declared differentially expressed if its adjusted p-value was lower than 0.05.

Hierarchical clustering of differentially expressed genes (DEGs) was performed using the MeV software (v4.9) with Pearson correlation and average linkage to identify SA187- and nitrate-responsive gene expression clusters. GO term enrichment analysis of DEG clusters was conducted using the R package clusterProfiler (Yu *et al*., 2012). Terms with adjusted p-values (FDR) < 0.05 were considered significant.

#### ii. qRT-PCR

cDNA synthesis was performed using the ImProm-II™ Reverse Transcription System (Promega) with oligo-dT primers. Quantitative real-time PCR was performed using SYBR Green I Master Mix (Roche) on a Bio-Rad CFX384 Real-Time PCR Detection System. Each reaction was run with technical triplicates. Gene expression was normalized to *ACTIN2* and *YSL8*, and relative expression levels were calculated using the ΔΔCt method (Livak & Schmittgen, 2001).

### H. In silico promoter analysis

Promoter sequences (2,000 bp upstream of the transcription start site, TSS) of Arabidopsis thaliana NRT2.5 (AT1G12940) and NRT2.6 (AT3G45060) were retrieved from TAIR10. Predicted transcription factor binding motifs were identified using the PlantPAN 4.0 platform (Chow et al., 2024) with default parameters. Putative binding sites for EIN3, EIL1, EIL2, and EIL3 transcription factors were mapped and visualized. Graphical promoter maps were generated using Microsoft PowerPoint for clarity.

### I. Statistical analyses and visualization

qRT-PCR data were analyzed using CFX Manager software version 2.1 (Bio-Rad). All other statistical analyses were performed in R (v4.2.1). Growth parameters (FW, PRL, and LRD) were compared using two-tailed unpaired Student’s *t*-tests. Elemental composition (C, N, and C:N ratio), nitrate contents and bacterial colonization datasets were analyzed using the non-parametric Mann–Whitney U test. Data visualization, including boxplots and bar graphs with error bars, was carried out using ggplot2, ggpubr, and rstatix packages in R. A threshold of *p* < 0.05 was used to determine statistical significance, with significance levels indicated directly on plots.

## III. Results

### A. Effect of SA187 on plant growth under varying nitrate conditions

To evaluate the impact of SA187 inoculation on seedling growth with respect to nitrate concentration, *Arabidopsis thaliana* seedlings were grown on solid media containing various concentrations of nitrate (20, 10, 2, 1, and 0.2 mM NO_3_^-^), with or without SA187 inoculation. Growth responses were quantified by measuring total FW (TFW), PRL, and LRD (Figure 1a–c). At high nitrate availability (20 mM), SA187 inoculation had no significant effect on TFW and PRL compared to mock controls (Figure 1a, b), however a significant increase in LRD (+27.2%) was observed (Figure 1c). As external nitrate concentration declined, SA187-mediated growth enhancements became progressively more pronounced. At 10 mM, SA187 treatment significantly increased TFW by 83.1%, PRL by 18.1%, and LRD by 32.9% (Figure 1a–c) and similar responses were seen at 2 mM nitrate, with SA187 increasing TFW by 90%, PRL by 13.4%, and LRD by 30.8%. At 1 mM nitrate, the beneficial effects of SA187 further intensified for TFW (+108.5%) and PRL (+29.3%), but the LRD increase (+32%) remained comparable to that observed at 2 and 10 mM (Figure 1a–c). Under the lowest nitrate supply (0.2 mM), the effect of SA187 inoculation was exceptionally pronounced, leading to increases of 295.5% (TFW), 121.6% (PRL), and 274.9% (LRD) compared to mock controls (Figure 1a–c). Taken together, these results indicate that SA187 substantially improves both shoot and root development, specifically when nitrate availability is low, with the magnitude of the beneficial effects inversely correlated with external nitrate levels.

**Figure 1.**
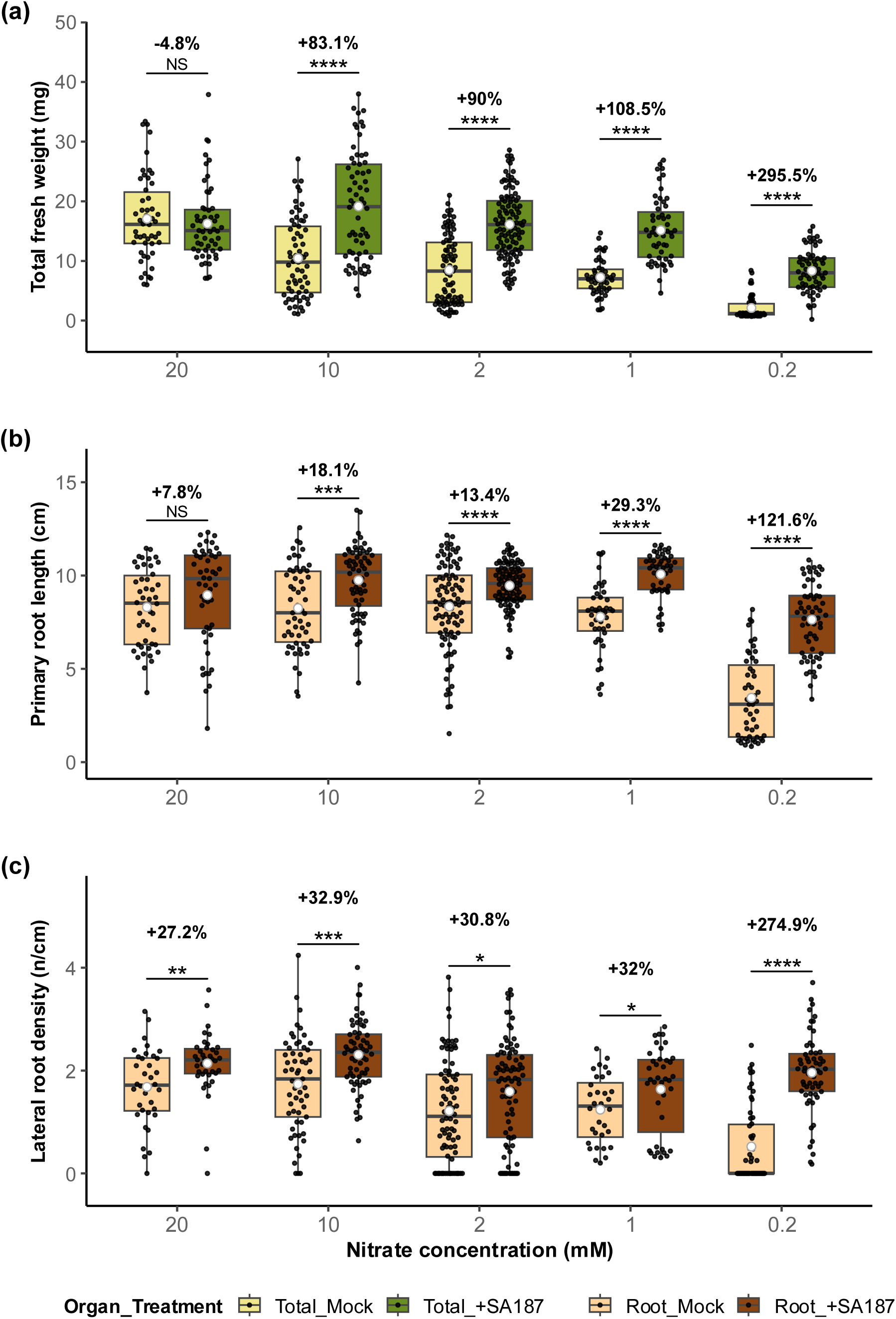
The beneficial effect of SA187 on *Arabidopsis thaliana* plant growth parameters under high and low nitrate. (a) Total fresh weight (mg) per seedling; (b) Primary root length (cm); (c) Lateral root density (n/cm = number/cm). Boxplots depict the median (horizontal line), interquartile range (box), and whiskers to 1.5× IQR; individual data points are overlaid as black dots, and white circles denote group means (n ≥ 40). Percentage above each bracket gives the change in mean with +SA187 relative to Mock. Statistical significance between treatments at each nitrate level was determined by two-sided, unpaired Student’s t-test: *P < 0.05; **P < 0.01; ***P < 0.001; ****P < 0.0001; NS, not significant.

### B. Effect of SA187 on C, N, and nitrate contents

To investigate how SA187 affected nitrate accumulation and internal nitrate partitioning, total C, total N, the C:N ratio, and nitrate content were measured in both shoots and roots of *Arabidopsis thaliana* grown under high (20 mM) and low (1 mM) nitrate conditions, with or without SA187 inoculation (Figure 2a–d). SA187 inoculation did not significantly alter shoot C content under either nitrate condition, but slightly reduced root C content at low nitrate (Figure 2a). N content exhibited tissue-specific responses: SA187 significantly increased shoot N content but decreased root N content under low nitrate conditions (Figure 2b). However, no significant changes in N content were observed under high nitrate conditions (20 mM) (Figure 2b). These variations in C and N content led to a significantly lower C:N ratio upon SA187 inoculation at 1 mM nitrate, both in shoots and in roots (Figure 2c). Most notably, SA187 induced a significant increase in nitrate content under low nitrate (1 mM) in both shoots and roots when compared to controls (Figure 2d). At high nitrate conditions (20 mM), on the other hand, SA187 had no significant effect on nitrate content. Collectively, these findings suggest the role of SA187 inoculation in improving nitrate accumulation, and shifting N-allocation and nitrate translocation towards aerial tissues under conditions of low nitrate. Such SA187-dependent changes could contribute to the observed increase in seedling growth.

**Figure 2.**
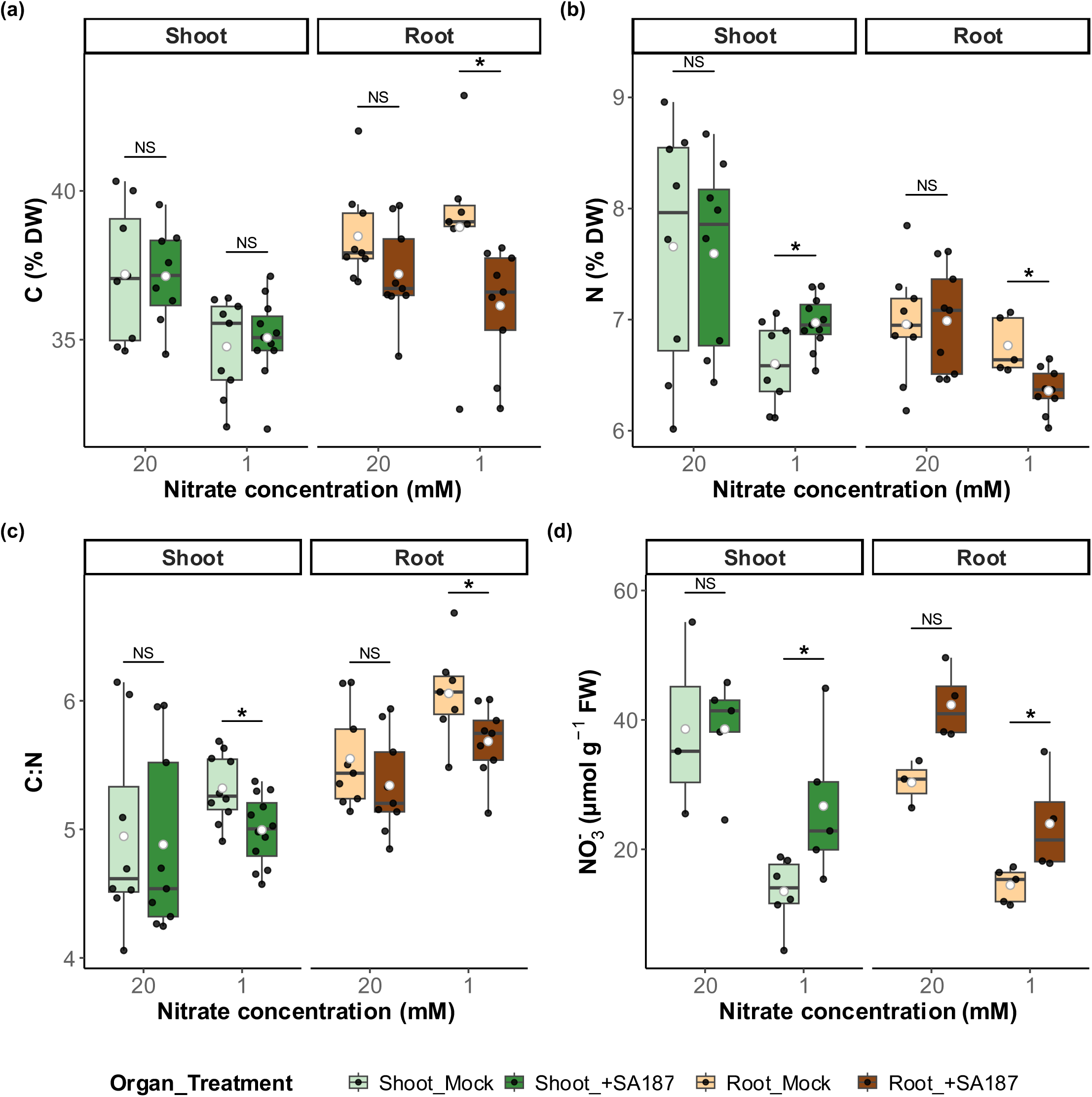
The effect of SA187 on *Arabidopsis thaliana* shoot and root C and N composition under high and low nitrate. Panels show boxplots of (a) total C content (% DW), (b) total N content (% DW), (c) C∶N ratio, and (d) nitrate content (µmol g ¹ FW) in shoots and roots for mock and SA187 treatments at each nitrate concentration. Boxplots display the median (center line), interquartile range (box), and whiskers (1.5× IQR); individual samples are plotted as black dots, white circles mark group means. Mock and +SA187 are distinguished by lighter versus darker colors. Wilcoxon rank sum test (Mann–Whitney U test) was used to compare treatments within each nitrate condition (significance: * = p < 0.05, ** = p < 0.01, *** = p < 0.001, NS = non-significant).

### C. SA187 mediated transcriptomic reprogramming of Arabidopsis shoots and roots

To gain insights into the molecular basis of SA187-mediated growth enhancement in the presence of low nitrate, transcriptome analyses (RNA-seq) were performed on Arabidopsis shoots and roots under high (20 mM) and low (1 mM) nitrate conditions with and without SA187 inoculation. Differential gene expression analysis identified 2,280 and 1,005 DEGs in shoots and roots, respectively (Supplementary Figure S1, Supplementary Tables S2 and S3). Hierarchical clustering and Gene Ontology (GO) enrichment analyses were conducted to categorize the DEGs based on their expression patterns and functional annotations.

In shoots, hierarchical clustering revealed distinct expression patterns. Cluster 4 (517 genes), the largest shoot cluster, showed significant gene expression induction by SA187 in both nitrate conditions, although predominantly under high nitrate, and included genes associated with defense, immune response, oxidative stress, ethylene, salicylic acid (SA) signaling and metabolic processes, jasmonic acid (JA), cell wall thickening and response to N-compounds. Cluster 10 (239 genes) exhibited a nitrate-dependent regulation, being upregulated under low nitrate alone but downregulated by SA187 treatment under low nitrate conditions. This cluster was enriched in genes linked to JA response, oxidative stress, amino acid metabolism, flavonoid biosynthesis, and glutathione metabolism. Genes in cluster 3 (81 genes) and cluster 9 (284 genes) demonstrated SA187-specific induction regardless of nitrate concentration. These clusters included genes involved in systemic resistance, response to N-compounds, lateral root development, camalexin biosynthesis and auxin and ethylene signaling with the induction of auxin related genes such as *ARGOS* (auxin-regulated gene involved in organ size), *IAA2 (*indole-3-acetic acid inducible 2), the Auxin-responsive GH3 family protein *BRU6* or ethylene responsive genre as *ERF1/2/5/15* (ethylene responsive factor 2), *EBP* (ethylene-responsive element binding protein) or *WRKY71* (Fig. 3a).

**Figure 3.**
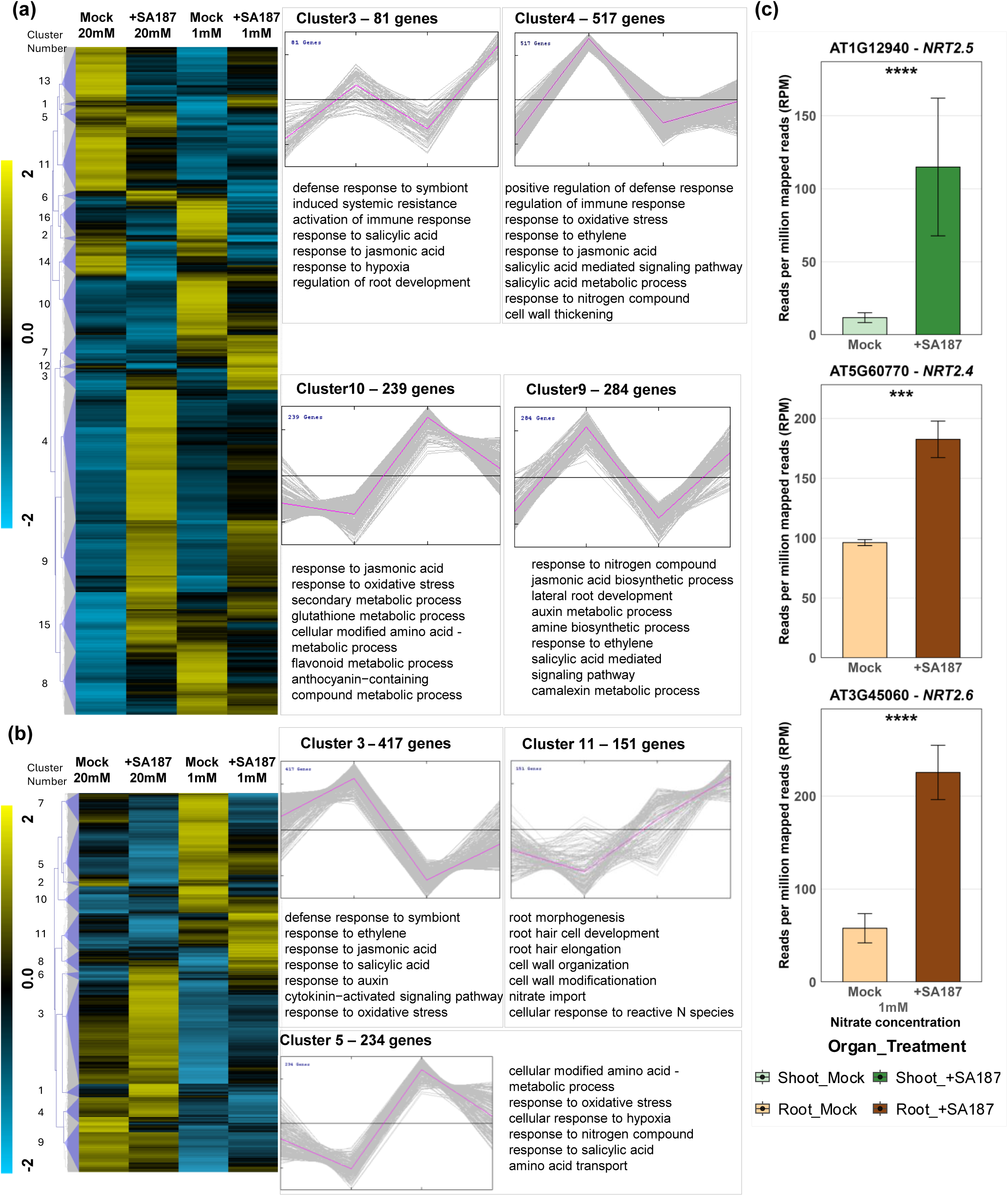
Global transcriptomic responses to SA187 in *Arabidopsis thaliana* shoots and roots under high and low nitrate. Hierarchical clustering heat maps of normalized gene expression (genes mean-centered across all samples) in **(a)** shoots and **(b)** roots of wild-type Col-0 seedlings treated with Mock or SA187 under high (20 mM) and low (1 mM) nitrate. Clustering was performed in MeV v4 using Pearson correlation and average linkage. Key clusters showing the most relevant induction or repression patterns are highlighted and annotated with enriched Gene Ontology (GO) biological processes obtained by G:Profiler. The color scale denotes relative expression (yellow = high, blue = low). The original mean counts were subjected to data adjustment by normalizing genes across all samples. Hierarchical clustering was performed using average linkage under Pearson correlation (MeV version 4). **(c)** Bar plots representing RNA-seq differential expression of selected HATS candidate genes (AT1G12940, *NRT2.5*; AT3G45060, *NRT2.6*; AT5G60770, *NRT2.4*) significantly regulated by SA187 treatment at 1 mM nitrate. Significance was determined by RNA-seq data analysis, with asterisks (***) denoting statistically significant differences (p-value < 0.001). (See also: Figure S1, Table S2 and Table S3)

Root transcriptome profiling also revealed distinct clusters of gene expression. Cluster 3 (417 genes), the largest root cluster, exhibited down-regulation under low nitrate but pronounced induction by SA187 across both nitrate conditions. Genes in this cluster were primarily related to defense, ethylene, JA, SA, auxin, cytokinin signaling, and oxidative stress responses. Cluster 5 (234 genes) showed an inverse expression pattern to cluster 3, being upregulated under low nitrate but strongly downregulated by SA187 at both nitrate concentrations. It included genes involved in amino acid transport and metabolism, N-signaling, and oxidative stress responses. Notably, cluster 11 (151 genes) was selectively induced by SA187 under low nitrate and enriched for genes involved in root morphogenesis, nitrate import, and responses to reactive N-species (Figure 3b), suggesting a condition-specific program optimized for nutrient accumulation under stress. Targeted examination of HATS highlighted a strong SA187-induced expression of *NRT2.5* (At1g12940) in shoots, and *NRT2.4* (At5g60770) and *NRT2.6* (At3g45060) in roots, under low nitrate conditions (Fig. 3c).

Together, these findings suggested that SA187 orchestrated a transcriptional network integrating nutrient transport, hormone signaling, stress buffering, and developmental remodeling to promote plant adaptation under a low nitrate condition.

### D. Role of high-affinity nitrate transporters NRT2.5, NRT2.6 and ethylene signaling in SA187-mediated growth promotion

To evaluate the functional requirement of ethylene signaling and specific HATS in SA187-induced plant growth benefits under low nitrate conditions, phenotypic analyses were performed using ethylene-insensitive (*ein2-1*), HATS single (*nrt2.5* , *nrt2.6*) and double (*nrt2.5 × nrt2.6*) mutants under low nitrate (1 mM). Wild-type plants (Col-0) exhibited substantial growth enhancement upon SA187 inoculation, as expected. However, none of the tested mutants displayed significant growth improvements upon SA187 inoculation, indicating that both nitrate transporters and a functional ethylene signaling were indispensable for the growth-promoting effects of SA187 under low nitrate (Figure 4a). Interestingly, in *ein2-1* and *nrt2.5*, SA187 slightly reduced growth under low nitrate, suggesting a potential antagonistic effect in the absence of functional ethylene signaling or NRT2.5.

**Figure 4.**
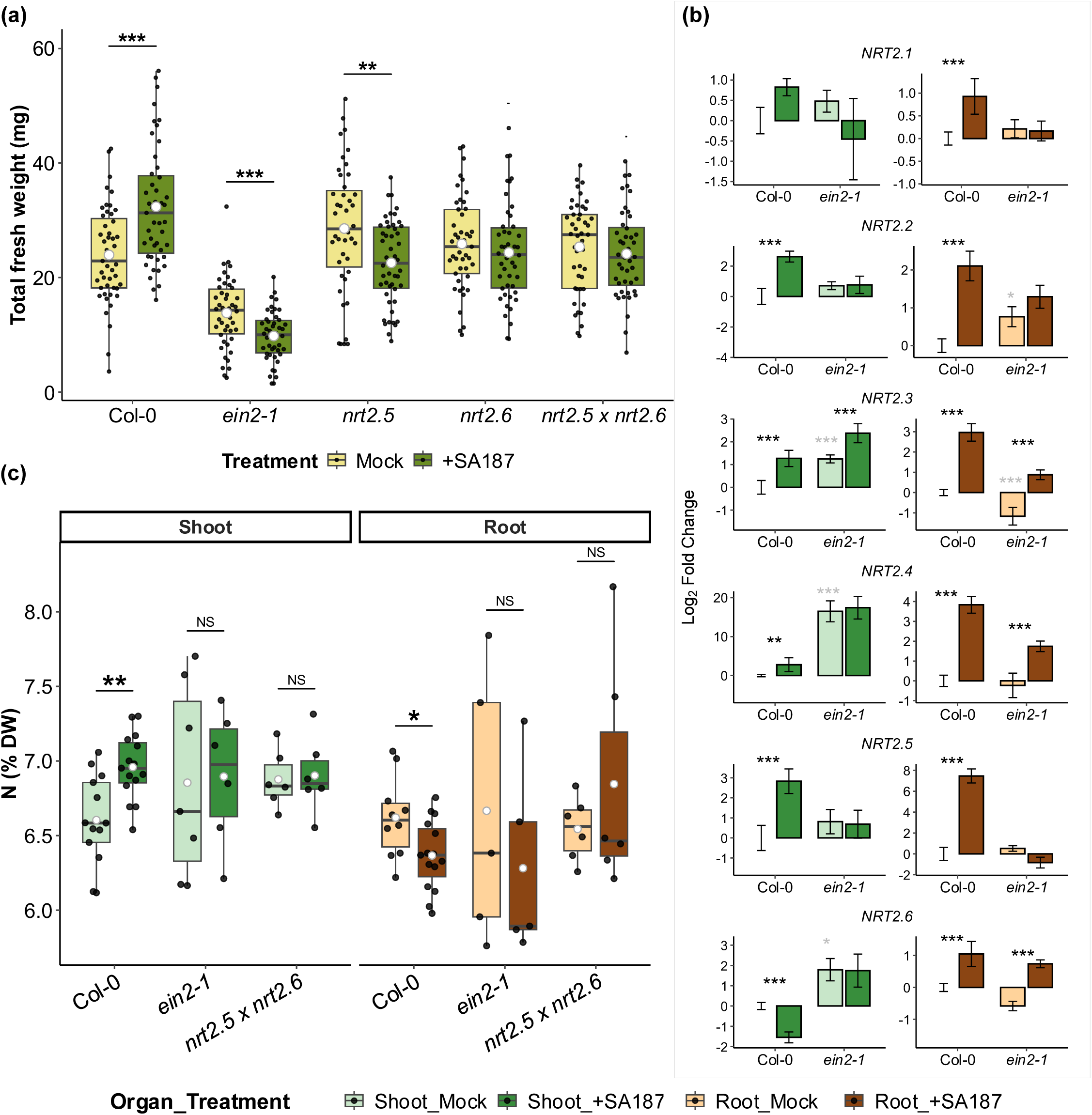
Effect of SA187 inoculation on *Arabidopsis thaliana* wild-type (Col-0) and mutant seedlings grown under low nitrate. (a) Total FW of seedlings grown with Mock or +SA187 treatments. Boxplots show median (center line), interquartile range (box) and whiskers to 1.5× IQR; individual plants are overlaid as black dots and white circles denote group means (n ≥ 40). Percentage above each bracket gives the relative change in mean fresh weight with +SA187 versus Mock. Statistical significance was determined by two-sided unpaired t-tests within each genotype (**P < 0.01, ***P < 0.001). (b) qRT-PCR analysis of high-affinity nitrate transporter genes in shoots (upper panels) and roots (lower panels) of Arabidopsis Col-0 and *ein2-1* seedlings. Expression was calculated by the ΔΔCt method and is shown as log fold-change relative to the Col-0 Mock samples for each tissue. Black asterisks indicate significance between Mock and +SA187 treatments (*P < 0.05; **P < 0.01; ***P < 0.001); grey asterisks denote differences between Col-0 and *ein2-1* under the mock treatment. Error bars represent ± SEM of three biological replicates (n ≥ 3). (c) Total N content (% DW) in shoots and roots under mock or SA187 treatments in Col-0, *ein2-1*, and *nrt2.5×nrt2.6* seedlings grown under 1 mM nitrate. Boxplots display the median (center line), interquartile range (box), and whiskers (1.5× IQR); individual samples are plotted as black dots, white circles mark group means (n ≥ 3). Mock and +SA187 are distinguished by lighter versus darker colored tones. Wilcoxon rank sum test (Mann–Whitney U test) was used to compare treatments within each nitrate condition (significance: * = p < 0.05, ** = p < 0.01, *** = p < 0.001).

SA187 colonized roots and shoots of Col-0, *ein2-1*, *nrt2.5*, and *nrt2.6* at comparable levels, confirming that impaired colonization is not the reason for the loss of growth promotion (Supplemental figure S3). To test whether the absence of SA187-mediated growth promotion in *ein2-1*, *nrt2.5*, and *nrt2.6* was specific to low nitrate, their response under 20 mM nitrate was also analyzed. Similar to Col-0, none of the mutants showed significant differences in seedling fresh weight upon inoculation, indicating that the lack of growth promotion is specific to low-nitrate conditions (Supplemental Fig. S4). This is not unexpected, since at such high nitrate availability plants likely reach their maximal growth potential, leaving little scope for additional growth promotion by SA187.

To further dissect the regulatory link between ethylene signaling and nitrate transporter expression, transcript levels of six HATS genes (*NRT2.1–NRT2.6*) were quantified by qRT-PCR in shoots and roots of Col-0 and *ein2-1* plants at 1mM nitrate, with and without SA187 inoculation (Figure 4b). NRT2.7 was excluded due to a consistently undetectable expression. In mock-inoculated *ein2-1* plants, baseline expression of *NRT2.3*, *NRT2.4*, and *NRT2.6* was higher in shoots compared to Col-0, while in roots, *NRT2.3* and *NRT2.6* were slightly downregulated, and *NRT2.2* was slightly upregulated. To examine how SA187 differentially modulated HATS expression in Col-0 and *ein2-1*, qRT-PCR analyses were compared following SA187 inoculation of both genotypes. In shoots, *NRT2.2*, *NRT2.3*, *NRT2.4*, and *NRT2.5* were upregulated by SA187 in Col-0. However, in *ein2-1*, SA187-mediated upregulation was only observed for *NRT2.3* in both organs and for *NRT2.4* in roots, suggesting a partial disruption of the regulatory response. Notably, *NRT2.6* was downregulated by SA187 in Col-0 shoots but showed no significant regulation in *ein2-1*. In roots, SA187 induced the expression of all tested HATS in Col-0. In contrast, SA187 failed to upregulate *NRT2.1*, *NRT2.2*, and *NRT2.5* in *ein2-1* roots, despite a normal induction of the other HATS. Amongst the examined HATS, *NRT2.5* exhibited the highest fold-change in response to SA187 in both organs when compared to mock controls, highlighting it as a potential major component in SA187-mediated growth enhancement under low nitrate conditions. Taken together, these results revealed that ethylene signaling was not essential for the basal expression of most HATS but it was critical for their SA187-induced activation. A major part of the SA187-induced modulation of HATS expression was observed predominantly in roots, although some notable effects were also detected in shoots. The *ein2-1* mutation specifically disrupted SA187-mediated induction of *NRT2.2* and *NRT2.5* in both shoots and roots, *NRT2.4* and *NRT2.6* in shoots, and *NRT2.1* in roots. These findings support a model in which SA187 leverages ethylene signaling to fine-tune the expression of a subset of nitrate transporters that improve seedling growth and nitrate content under low nitrate conditions.

### E. Importance of EIN2, NRT2.5, and NRT2.6 in SA187-induced nitrogen accumulation

To determine whether the SA187-induced upregulation of HATS and growth promotion is associated with changes in N-allocation, total N content (% DW) was measured in shoots and roots of Col-0, *ein2-1*, and the *nrt2.5×nrt2.6* double mutant under low nitrate conditions (1 mM NO_3_^-^) with and without SA187 inoculation (Figure 4c). In wild-type (Col-0) plants, SA187 inoculation significantly increased shoot N-content compared to mock controls. However, this beneficial effect was abolished in both *ein2-1* and *nrt2.5×nrt2.6* shoots, with no significant differences detected between inoculated and mock-treated plants (Fig. 4c). In roots, SA187 did not significantly alter N-content in wild-type and mutant lines, suggesting that SA187 impacted N-allocation to the shoot under these experimental conditions (Fig. 4c). These results further confirmed the involvement of ethylene signaling, *NRT2.5*, and *NRT2.6* in SA187-induced promotion of shoot N content.

### F. EIN3 is required for SA187-mediated growth promotion and NRT2.5/2.6 harbor EIN3/EIL binding motifs

To determine whether SA187-mediated growth promotion requires downstream components of ethylene signaling, *ein3-1* mutants were analyzed under low nitrate (1 mM NO) with or without SA187 inoculation. While SA187 significantly increased FW in Col-0 (+93.3%), no growth enhancement was observed in *ein3-1*, confirming that EIN3 is essential for the SA187-mediated growth response (Figure 5a).

**Figure 5.**
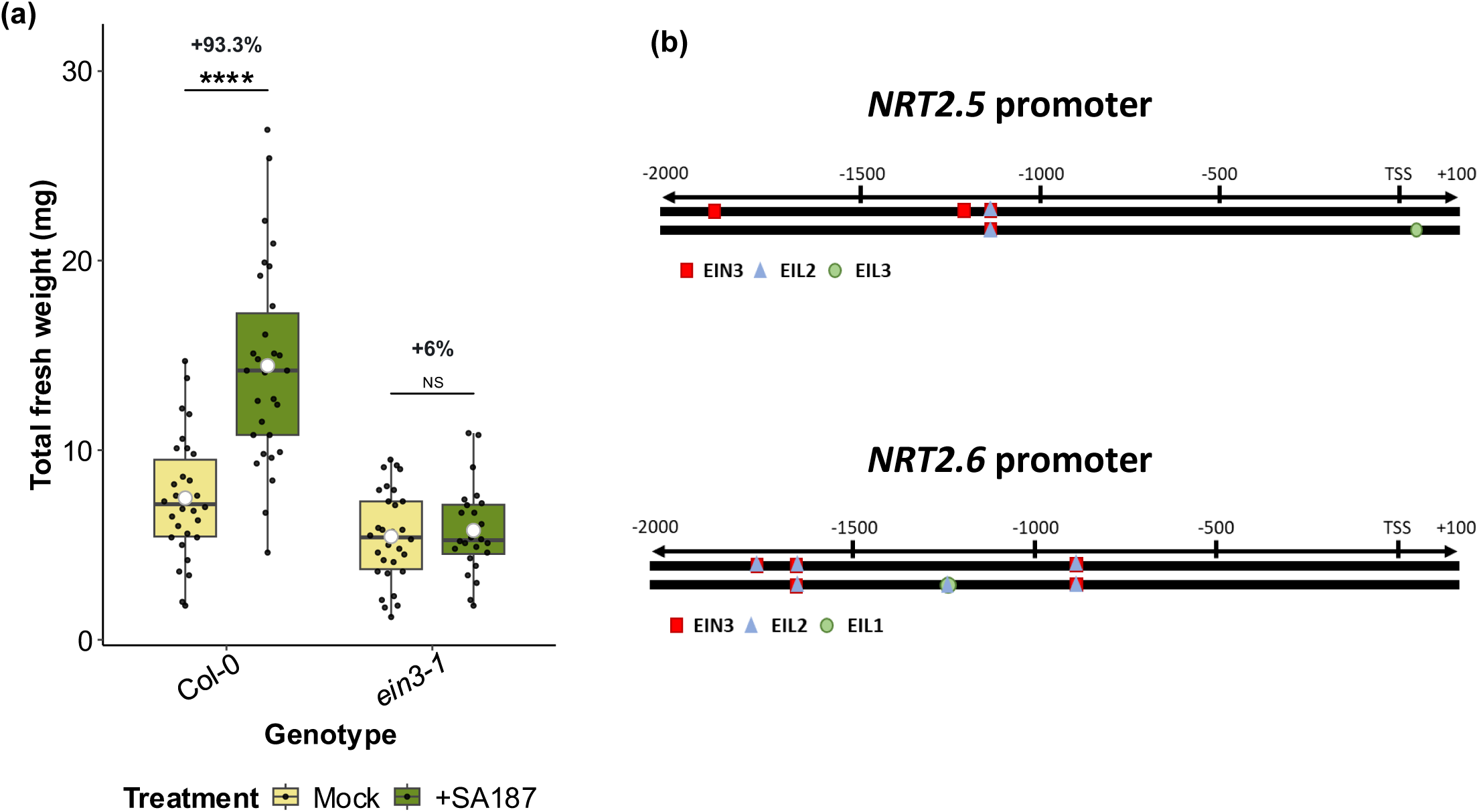
Simplified model of the regulatory influence of SA187 on the expression of *Arabidopsis thaliana* high-affinity nitrate transporter genes in shoots and roots. SA187 inoculation promotes plant growth under low nitrate by modulating the expression of HATS through both ethylene-dependent and independent pathways. *NRT2.5* is strongly upregulated by SA187 in an EIN2-dependent manner in both shoots and roots, while *NRT2.6* shows tissue-specific regulation: downregulated in shoots and upregulated in roots, with a partial ethylene independence. Solid arrows indicate confirmed regulatory relationships; dashed arrows denote hypothesized or indirect links. The model proposes that SA187 enhances nitrate uptake and aerial N-allocation via NRT2.5 and NRT2.6 thus contributing to improved growth under limited nitrate conditions. Supplementary Figure 1. Differential gene expression in *Arabidopsis thaliana* under high or low nitrate and with or without SA187 inoculation. Transcriptomic comparisons of Arabidopsis seedlings inoculated with SA187 versus mock-treated controls under low (1 mM) and high (20 mM) nitrate concentrations. (a) DEGs in shoot tissues. (b) DEGs in root tissues. Each comparison indicates the number of significantly upregulated (red) and downregulated (purple) genes identified by pairwise contrasts. Arrows represent the direction of each comparison.

**Figure 6.**
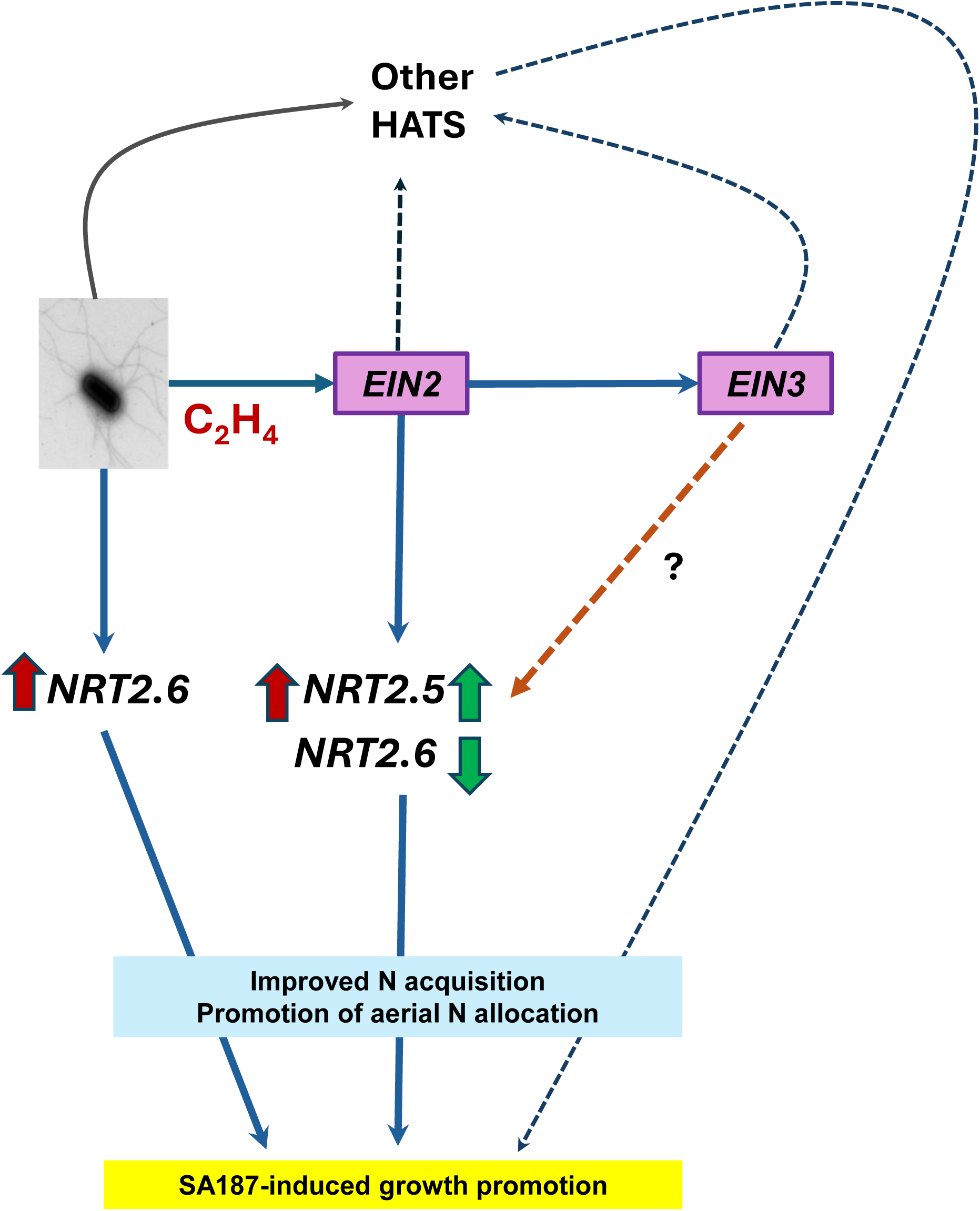
Simplified model of the regulatory influence of SA187 on the expression of Arabidopsis thaliana high-affinity nitrate transporter genes in shoots and roots. SA187 inoculation promotes plant growth under low nitrate by modulating the expression of HATS through both ethylene-dependent and independent pathways. *NRT2.5* is strongly upregulated by SA187 in an EIN2-dependent manner in both shoots and roots, while *NRT2.6* shows tissue-specific regulation: downregulated in shoots and upregulated in roots, with a partial ethylene independence. Solid arrows indicate confirmed regulatory relationships; dashed arrows denote hypothesized or indirect links. The model proposes that SA187 enhances nitrate accumulation and aerial N-allocation via *NRT2.5* and *NRT2.6* thus contributing to improved growth under limited nitrate conditions.

To explore whether NRT2.5 and NRT2.6 might be directly regulated by EIN3/EIL transcription factors, *in silico* promoter analyses were performed using PlantPAN 4.0. Both NRT2.5 and NRT2.6 promoters contained multiple predicted EIN3/EIL family binding sites within 2 kb upstream of the transcription start site (TSS) (Figure 5b). These motifs suggest potential direct transcriptional regulation by EIN3/EILs, linking ethylene signaling with the activation of nitrate transporter genes under SA187 inoculation.

## IV. Discussion

### A. SA187 enhances plant growth under low nitrate conditions

This study demonstrates that SA187 significantly promoted Arabidopsis growth under low nitrate conditions through a coordinated enhancement of seedling growth and root development, thus highlighting a potential role in limiting N-based fertilizer use. The observed growth-promoting effect was tightly linked to external nitrate availability. While only a negligible SA187 effect was seen under high nitrate (20 mM), SA187 significantly increased total FW, PRL, and LRD under low nitrate regimes (between 0.2–2 mM nitrate) (Figure 1). These findings position SA187 as a context-dependent growth promoting inoculant that becomes functionally more advantageous under low nitrate conditions, a hallmark of beneficial plant–microbe interactions to support host fitness under suboptimal environmental conditions. Indeed, previous studies have shown that SA187 does not promote plant growth under non-stress conditions but significantly enhances biomass and root system development under abiotic stresses such as salinity, heat, and high light stress (Andrés-Barrao et al., 2017; Andres-Barrao et al., 2021; de Zélicourt et al., 2018; Rolli et al., 2022; Shekhawat et al., 2024). Absence of growth promotion at higher N levels could also imply that uninoculated plants already reached maximal growth capacity under these conditions.

Interestingly, unlike many PGPB reports (Spaepen et al., 2008; Vacheron et al., 2013), SA187 did not reduce PRL, even though LRD was consistently increased across all nitrate concentrations, including 20 mM. This suggests that SA187 triggers a constitutive lateral root response without constraining primary root elongation. Moreover, although SA187 clearly enhanced LRD, our data cannot distinguish whether this effect reflects increased initiation of lateral roots or enhanced activation of pre-formed primordia. This distinction will help understanding root growth promotion mechanisms by SA187 but requires future dedicated studies.

Interestingly, SA187 consistently increased LRD across all tested nitrate concentrations including 20 mM when shoot biomass and PRL were unaffected. This constitutive increase in lateral root formation, regardless of nitrate status, suggests that SA187 may activate a basal root developmental response. This may improve the plant’s capacity to explore heterogeneous soil environments and exploit transient nutrient patches, an adaptive advantage even when nitrate supply is sufficient (Garnett *et al*., 2009). Enhancement of lateral root density has been observed previously in other PGPB interactions, where changes in root architecture were found to be driven by microbial modulation of plant hormone pathways, including auxins, cytokinins, and ethylene (Spaepen et al., 2008; Vacheron et al., 2013; Chen et al., 2024). The SA187 genome contains *tnaA*, encoding a tryptophanase that converts tryptophan into indole, a known auxin precursor, as well as an indole efflux pump (*acrEF*) (Andrés-Barrao *et al*., 2017; Synek *et al*., 2021). Indole and its derivatives contribute to the biosynthesis of indole-3-acetic acid (IAA) and indole-3-butyric acid (IBA), key auxins involved in lateral root initiation and elongation (Trujillo-Hernandez *et al*., 2020; Sun *et al*., 2023). It is therefore possible that SA187 influences host auxin dynamics either by directly producing auxin-like metabolites or by stimulation of host endogenous auxin.

The observed root remodeling may be underpinned by an additional layer of mutualism. It has been proposed that SA187 preferentially enters plant tissues via natural openings at lateral root emergence sites (Synek *et al*., 2021). From this perspective, increased LRD not only benefits the plant by expanding its nutrient absorption capacity but it will also facilitate bacterial entry and endophytic establishment. Such a dual-purpose modulation, benefiting both microbe and host, exemplifies the mutualistic sophistication of beneficial endophytes. Collectively, these observations are consistent with SA187-associated changes in root architecture and growth that could favor N accumulation under low nitrate, together with transcriptional regulation of high-affinity nitrate transporters; however, our dataset does not distinguish between these possibilities. accumulation

### B. SA187 impacts C and N content and improves nitrate accumulation under low nitrate conditions

SA187 inoculation enhanced Arabidopsis nitrate content and elemental analyses revealed an altered internal N-distribution under low nitrate conditions. This was seen especially under low-nitrate where SA187 increased shoot total N-content while either reducing (Fig. 2b, Fig. 4c)in the root system. This was accompanied by elevated nitrate levels in both organs in SA187 inoculated seedlings compared to wild-type Col-0. This suggested that SA187 not only facilitated root nitrate uptake but also enhanced its translocation to aerial tissues. Therefore, it appeared that SA187 mediated an altered N-allocation that favored N investment in aerial parts compared to roots. This was only observed under low nitrate, when sustaining photosynthetic efficiency and shoot biomass depends heavily on optimal N-partitioning. The observed shift in N-content from roots to shoots could reflect a strategy to redirect resources to maximize growth under suboptimal N-conditions. Improvement in both nitrate uptake, improvement if overall N status and accumulation of shoot N content has been demonstrated previously in multiple plant-microbe systems. For example, *Achromobacter* sp. 3-17 inoculation significantly improved biomass production and doubled nitrate uptakeratein *Brassica napus* (Bertrand *et al*., 2000). Likewise, volatile organic compounds from *Bacillus velezensis* SQR9 promoted uptake rate of different forms of nitrogen, total nitrogen content and growth in both Arabidopsis and rice (Chen *et al*., 2024). Another *Enterobacter* species, *E. cloacae* subsp. *dissolvens*, was shown to improve N-content and yield of teff varieties under field conditions, indicating the broader potential of this genus in improving plant N-nutrition (Tsegaye *et al*., 2022). However, contrasting results have also been reported where PGPB improved plant growth without proportionate increases in nitrate accumulation or uptake. For example, *Phyllobacterium brassicacearum* STM196 enhanced shoot total N content at all nitrate supplies (0.5–10 mM NO) but increased nitrate pools only at 0.5 mM. At higher concentrations, nitrate pools remained unchanged or declined, even though shoot nitrogen accumulation continued to rise. Transient increases in nitrate influx were detected after 24 h of inoculation but disappeared after 8 days, indicating dynamic, concentration-dependent regulation (*Mantelin et al.*, 2005). Collectively, studies show that PGPB-induced enhancement of plant nitrogen status depends on bacterial strain, nitrate concentration, and nitrogen form, rather than representing a universally fixed outcome and most of the studies have reported correlative outcomes than mechanistic causation.

Beyond uptake, the effective transport and assimilation of nitrate into amino acids and proteins is a key determinant of NUE and overall plant growth and productivity. By enhancing both nitrate uptake and its allocation to shoots, PGPB may help plants optimize N-use under stress conditions. Upregulated expression of nitrate transporters involved in xylem loading, such as AtNRT1.5, has been reported in response to beneficial microbes, facilitating enhanced nitrate flow to shoot tissues (Calvo *et al*., 2019). Such enhanced translocation may not only support vegetative growth but also influence seed development and nutrient remobilization during reproductive stages. Indeed, microbial inoculation has been shown to improve N-remobilization efficiency in crop plants, benefiting grain protein content and yield (Adesemoye *et al*., 2008). Although the current study focused on vegetative development, it is plausible that SA187-mediated enhancements in nitrate uptake and N-allocation may also support reproductive success and seed filling, contributing to overall NUE, a key trait for achieving sustainable crop productivity under N-limited conditions (Congreves *et al*., 2021).

The C/N ratio in plant tissues is an indicator of metabolic balance and nutritional status. Efficient N-assimilation requires the incorporation of C-skeletons into amino acids and structural components, thereby limiting the buildup of unused C-rich metabolites when N is scarce (Elser *et al*., 2010; Baslam *et al*., 2020; Li *et al*., 2022). In line with this, C:N ratios were elevated under low nitrate availability. However, SA187 inoculation significantly reduced C:N ratios under these conditions, suggesting improved N accumulation and assimilation. Notably, SA187-inoculated plants under low nitrate exhibited shoot C:N ratios comparable to those observed in plants grown under a high nitrate supply. This suggests that SA187 enhanced nitrate accumulation sufficiently to restore metabolic balance even in low-nitrate environments. From an agronomic perspective, plants with low C:N ratios often have high protein contents, improved nutritional quality, and produce residues that decompose more readily, enhancing soil fertility and nutrient recycling (Hertzler *et al*., 2020; De Silva *et al*., 2023).

Collectively, these results point to a dual contribution of SA187 in facilitating nitrate uptake and optimizing its internal use and allocation. Such a capacity highlights the potential of SA187 as a bioinoculant for improving plant N-status, especially in the context of reducing the use of synthetic fertilizers while maintaining the productivity of low-input agricultural systems.

### C. Transcriptomic analysis reveals SA187-induced reprogramming of nitrogen and hormone signaling pathways under low nitrogen

Transcriptomic profiling revealed that SA187 induced a coordinated and condition-specific reprogramming of *Arabidopsis thaliana* gene expression (Figure 3). This reprogramming encompassed multiple layers of regulatory networks including nutrient accumulation, hormone-mediated development, immune modulation, and cellular stress responses. The observed expression patterns reflected a potential dynamic microbial modulation of host physiology to improve growth and resilience in low nitrate environments.

The notable induction of defense-and immune-related genes by SA187, especially under sufficient nitrate conditions, suggested an underlying mechanism of induced systemic resistance (ISR). It is a common plant defense strategy triggered by beneficial microbes, allowing plants to rapidly and robustly respond to eventual biotic stress conditions (Van der Ent *et al*., 2009; Pieterse *et al*., 2014). This priming effect by SA187 aligns closely with previous reports demonstrating ISR induction by beneficial rhizobacteria (Zamioudis et al., 2013; de Zélicourt et al., 2018). The largest SA187-induced gene cluster in shoots (Cluster 4) was enriched for genes associated with defense signaling, oxidative stress responses, and hormone pathways including ethylene, SA and JA. These genes were strongly induced by SA187 under high nitrate conditions and to a lesser extent under low nitrate, suggesting a nitrate-independent component of defense priming. Such transcriptional activation is consistent with microbe-ISR, a process where non-pathogenic rhizobacteria enhance plant immunity (Zehnder *et al*., 2001; Choudhary *et al*., 2007; Van der Ent *et al*., 2009; Pieterse *et al*., 2014). The concurrent induction of SA and JA signaling components points to a nuanced immune regulation, potentially enabling both basal defense and biotic stress tolerance. Studies have shown that an increased N-supply can lead to plant susceptibility to pathogens (Mur *et al*., 2016; Jeon, 2019; Maina *et al*., 2024). Interestingly, although SA187 did not enhance biomass under high nitrate, its activation of immune pathways suggested it could confer fitness advantages by strengthening plant immunity. Consistent with this idea, shoot Clusters 3 and 9 displayed SA187-induced expression irrespective of N-status. These clusters were enriched in genes responding to microbial symbionts, hormone signaling (SA, JA, auxin, ethylene), N-compounds, and root developmental regulators such as lateral root initiation and camalexin biosynthesis. The nitrate-independent induction of these clusters suggested a basal SA187-triggered transcriptional program integrating developmental cues with immune modulation, likely enabling dual growth-promotion and stress-buffering effects. The co-enrichment of hypoxia-responsive genes could be linked to bacterial modulation of the rhizosphere redox environment, potentially altering local oxygen tension or reactive oxygen species homeostasis, both known to influence root development and immune status (Bailey-Serres & Voesenek, 2008; Considine & Foyer, 2014).

On the other hand, shoot Cluster 10 displayed a nitrate-sensitive signature where genes were upregulated under low nitrate but repressed upon SA187 inoculation. This cluster included genes related to detoxification, glutathione metabolism enzymes, and flavonoid biosynthetic, classical metabolic markers of cellular stress responses to N-deprivation and oxidative imbalance (Lea *et al*., 2007; Krapp *et al*., 2011; Curci *et al*., 2017; Liang & He, 2018; Li *et al*., 2023; Yang *et al*., 2024). SA187-mediated suppression of these stress markers under low nitrate again suggested a buffering effect, whereby SA187 reduced the perceived severity or molecular consequences of low nitrate.

Root transcriptome profiling revealed a comparable complexity. The largest cluster (Cluster 3) was strongly induced by SA187 under both nitrate regimes and contained genes involved in immune signaling, phytohormone pathways (ethylene, SA, JA, auxin, cytokinin), and oxidative stress responses. SA187 is known to colonize the root endosphere, therefore this widespread induction probably reflected direct microbe-host signaling at the root interface. Cluster 5 was characterized by a classical stress signature under low nitrate that was downregulated by SA187. This included genes related to amino acid metabolism, redox balance, and hypoxia, consistent with earlier findings that SA187 alleviates cellular stress via metabolic reprogramming (de Zélicourt *et al*., 2018; Andres-Barrao *et al*., 2021; Shekhawat *et al*., 2024). A strikingly distinct pattern was observed in Cluster 11, where genes were upregulated by SA187 only under low nitrate. Functional enrichment revealed root morphogenesis, root hair elongation, nitrate import, and cellular responses to reactive N species. The exclusive upregulation of this cluster under SA187–low nitrate conditions suggested a tightly controlled transcriptional module integrating nutrient signals with bacterial cues to selectively activate root developmental pathways when most needed.

It is interesting to note that, while SA187 is preferentially located in root, shoot transcriptome displays the most abundant DEG in response to SA187 inoculation compared to root transcriptome (Supplemental Figure S1), a feature already highlighted in a previous study (Andrés-Barrao et al., 2021). This might reflect the involvement of systemic signaling pathways, that will largely affect development and metabolism, thanks to root-to-shoot communication signals such as cytokinin as identified in the root transcriptome, but also the possible participation of small signaling peptides or microRNAs (Shanks et al., 2024).

Among the most functionally relevant findings was the upregulation of high-affinity nitrate transporter genes (*NRT2.5* in shoots, *NRT2.4* and *NRT2.6* in roots). These transporters are critical under low external nitrate concentrations (Lezhneva et al., 2014; Krapp et al., 2014). Their induction under low nitrate conditions suggested that SA187 benefits observed in terms of growth and nitrate accumulation and partitioning concerned NRT2.4, NRT2.5 and NRT2.6.

Collectively, the transcriptomic data supported a model in which SA187 dynamically rewired plant gene expression to coordinate developmental and metabolic adaptations, enhance nitrate accumulation and allocation, and buffer physiological stress responses. These multifaceted changes may enable SA187-inoculated plants to sustain growth and development under low nitrate conditions and position this endophyte as a potent microbial tool for improving NUE and resilience in crops.

### D. SA187-mediated growth promotion requires coordinated action of ethylene signaling and high-affinity nitrate transporters

Focusing on ethylene signaling and on NRT2.5 and NRT2.6 was done because SA187-mediated growth benefits have previously been linked to ethylene (de Zélicourt et al., 2018), and these transporters were shown by Kechid et al. (2013) to be essential for PGPB-induced growth promotion under low nitrogen. Functional analyses suggested that both ethylene signaling and specific high-affinity nitrate transporters, particularly *NRT2.5* and *NRT2.6*, were required for SA187-mediated growth promotion under low nitrate conditions. Phenotypic analyses confirmed that the *ein2-1*, *nrt2.5*, *nrt2.6*, *nrt2.5×nrt2.6* mutants failed to show significant growth improvements following SA187 inoculation, unlike wild-type plants.

Indeed, previous reports have demonstrated the crucial regulatory role of ethylene in modulating nitrate transporter expression and root system architecture under varying N-conditions. Ethylene signaling, mediated through *EIN2/EIN3* pathways, has been previously shown to regulate critical nitrate transporter genes such as *NRT1.1* and *NRT2.1*, fine-tuning nitrate accumulation and root developmental responses to external N-availability (Tian *et al*., 2009; Zheng *et al*., 2013). For instance, ethylene-mediated repression of lateral root growth under high nitrate conditions required intact *EIN2-1* signaling, and ethylene-insensitive mutants (*ein2-1*, *etr1-3*) exhibited dysregulated root responses to nitrate signals (Tian *et al*., 2009). Additionally, Zheng et al. (2013) identified a feedback regulatory loop whereby low nitrate availability induced ethylene biosynthesis, subsequently repressing *NRT2.1* expression to avoid excessive nitrate uptake under prolonged deficiency. Under abiotic stresses, ethylene also modulated stress-induced nitrate allocation to roots via coordinated regulation of *NRT1.5* (xylem loading) and *NRT1.8* (xylem unloading), further exemplifying its central role in plant nutrient homeostasis and stress adaptation (Zhang *et al*., 2014).

The requirement for ethylene signaling in this context was further supported by SA187-induced expression of *NRT2.1–NRT2.6* in wild-type roots, and selective induction of *NRT2.2*, *NRT2.3*, *NRT2.4*, and *NRT2.5* in shoots. In contrast, *NRT2.6* was downregulated in shoots following SA187 treatment. Among all transporters, *NRT2.5* displayed the strongest induction, suggesting a central role in SA187-mediated enhancement of nitrate content and N-allocation. This transcriptional activation was largely abolished in *ein2-1*, which failed to upregulate key transporters including *NRT2.1* in roots, and *NRT2.2* and *NRT2.5* in both shoots and roots. These results clearly position these genes downstream of ethylene signaling in the SA187 response. Interestingly, SA187-induced *NRT2.6* downregulation in shoots was not observed in *ein2-1*, while root expression of *NRT2.6* remained upregulated in both genotypes, suggesting a partially ethylene-dependent regulatory pattern that was dependent in shoots but independent in roots.

The significance of *NRT2.5* in this context is consistent with its critical role under prolonged N-starvation. Indeed, *NRT2.5* was found to be essential not only for efficient nitrate uptake but also for phloem loading and N-remobilization, supporting plant growth under sustained N-limitation (Lezhneva *et al*., 2014). Similarly, *NRT2.4*, which was also significantly upregulated by SA187 in both organs, has been shown to contribute to nitrate uptake and remobilization, particularly in lateral roots under severe N-deficiency (Kiba *et al*., 2012). While a precise role for *NRT2.6* in nitrate transport remains poorly defined, earlier studies suggested its involvement in pathogen-induced oxidative stress responses rather than direct nitrate accumulation (Dechorgnat *et al*., 2012). This raises the possibility that its contribution to growth promotion extends beyond nitrate transport per se and may reflect regulatory or signaling functions.

*NRT2.1* was also differentially regulated, however its function was not pursue in detail because the *nrt2.1* mutant grows very poorly under our low nitrate (1mM), making it technically unfeasible to reliably characterize it further. Nevertheless, the known ethylene–NRT2.1 feedback loop (Zheng et al., 2013), suggests that NRT2.1 could intersect with SA187-triggered ethylene signaling. Resolving this question will require future dedicated analyses under conditions that permit *nrt2.1* plants to reach sufficient growth for reliable measurements.

Importantly, our findings confirming the requirement of *NRT2.5* and *NRT2.6* for SA187-mediated growth enhancement extend the earlier findings by Kechid et al. (2013) that *NRT2.5* and *NRT2.6* are also indispensable for growth promotion under low nitrogen by *Phyllobacterium brassicacearum* STM196, a phylogenetically distant and exclusively rhizobacterium. This parallel strongly suggests that the requirement of *NRT2.5* and *NRT2.6* represents a general feature of beneficial PGPB–plant interactions under low nitrogen, rather than a mechanism linked to the bacterial lifestyle.

The existence of both ethylene-dependent and ethylene-independent regulations of nitrate transporter expression by SA187 inoculation was observed when comparing wild-type and *ein2-1* seedlings. While *NRT2.2*, *NRT2.5*, *NRT2.6* (shoots) and *NRT2.1* (roots) clearly required ethylene signaling for SA187-induced expression, the regulation of *NRT2.3* in both organs and *NRT2.4* and *NRT2.6* in roots appeared to be independent of ethylene. This dual regulatory system may provide robustness to nitrate accumulation and translocation strategies under fluctuating environmental conditions where ethylene signaling may be perturbed, for example, under combined stress scenarios. Such regulatory flexibility shows the complexity of microbial modulation of plant nutrient responses and highlights the sophisticated molecular interplay microbial signals, hormone-mediated transcriptional networks, and nitrate transporter systems. While experimental evidences demonstrate that SA187-induced expression of NRT2.5 and NRT2.6 requires intact ethylene signaling, the regulatory connection remains correlative. Our *in-silico* analyses suggest potential EIN3 recognition sites upstream of NRT2.5, and NRT2.6 (Figure 5b, supplemental file W1). Alternatively, regulation may occur indirectly through EIN3-like (EIL) transcription factors acting downstream of EIN2. While these predictions remain to be experimentally validated through chromatin immunoprecipitation (ChIP) or reporter assays, they reinforce the hypothesis that EIN3-mediated transcriptional control may directly coordinate the activation of nitrate transporters under low-nitrogen conditions. This offers a plausible molecular connection between ethylene signaling and enhanced N allocation during SA187-mediated growth promotion.

Finally, although the exact trigger of ethylene signaling by SA187 remains unresolved in the scope of our work, previous studies showed that SA187 lacks ACC deaminase but produces 2-keto-4-methylthiobutyric acid (KMBA), which can be oxidized in planta to ethylene (de Zélicourt et al., 2018). This KMBA-dependent route provides a likely explanation for the activation of EIN2/EIN3 signaling observed here and is consistent with earlier demonstrations of SA187-induced ethylene responses under stress conditions.

### E. Ethylene signaling and nitrate transporters coordinate SA187-induced nitrogen allocation to shoots

The loss of SA187-mediated growth promotion in *ein2-1* and *nrt2.5×nrt2.6* mutants, combined with the altered expression of HATS in ethylene-insensitive *ein2-1*, led to the hypothesis that both ethylene signaling and functional NRT2.5 and NRT2.6 were critical for SA187-driven N-allocation to shoots. In wild-type plants, SA187 significantly increased shoot N-content, consistent with enhanced N status and preferential allocation to aerial tissues. The complete loss of this effect in *ein2-1* and *nrt2.5×nrt2.6* confirmed that both ethylene signaling and (at least one of) the HATS were required for this shift in N-allocation induced by SA187. The loss of this response in *ein2-1* reinforced the idea of ethylene as a central regulator in SA187-induced nitrate transport and N-allocation. Transcriptomic and qRT-PCR data consistently showed that *NRT2.5* expression was strongly induced by SA187 in wild-type plants but not in *ein2-1*, demonstrating that this transporter functioned downstream of ethylene signaling. These findings align with earlier studies indicating that ethylene can regulate nitrate transporter expression under N-limitation (Tian *et al*., 2009; Zheng *et al*., 2013), and support the view that ethylene functions as a key signaling hub linking microbial perception with nitrate uptake pathways. Similarly, the failure of SA187 to enhance shoot N-content in the *nrt2.5×nrt2.6* double mutant confirmed the essential contribution of these transporters to nitrate accumulation and distribution. SA187-dependent regulation of *NRT2.5* and *NRT2.6* under low nitrate, as observed in wild-type plants, indicates their roles in mediating improved N-allocation to shoots. Given that shoot N-accumulation correlated with growth improvement, this transporter-dependent redistribution is likely a prerequisite for the observed biomass gains.

Although SA187 inoculation coincided with higher shoot nitrogen content and induction of *NRT2.5* and *NRT2.6*, our data cannot distinguish whether this reflects direct changes in nitrate uptake rate, growth-driven increases in nutrient demand, or a combination of both. This limitation is compounded by the absence of separate shoot and root biomass measurements, total lateral root length, and direct nitrate flux assays. As a result, at least three scenarios, which are mutually non-exclusive, remain plausible: (i) SA187 primarily enhances root growth, indirectly improving shoot nutrition; (ii) SA187 stimulates shoot growth, thereby increasing nitrogen demand and accumulation; or (iii) SA187 directly modulates nitrate transporters, leading to improved nitrogen uptake and growth. Our results cannot discriminate between these alternatives, and future studies incorporating organ-specific biomass, detailed root architectural traits, and flux assays will be required to establish a causal effect of these mechanisms.

According to current physiological knowledge, nitrate delivery to aerial tissues depends primarily on root accumulation and subsequent xylem loading, rather than on shoot-localized high-affinity transporters. Thus, the requirement of NRT2.5 and NRT2.6 in SA187-mediated growth promotion cannot be explained by a canonical shoot transport function. Instead, their induction in shoots may reflect signaling roles that integrate nutrient status with hormonal and stress responses. Our findings also support the hypothesis that regulation of these transporters could influence nitrogen translocation to shoots, but this possibility remains to be rigorously tested, and these scenarios are not mutually exclusive. Distinguishing root-versus shoot-specific contributions of NRT2.5 and NRT2.6 will require targeted approaches; in principle, grafting experiments could provide definitive insights, although such methods remain technically challenging in Arabidopsis and will need to be pursued in future work.

Although ethylene emerged as the central regulator of the SA187 response, transcriptomic data also point to roles for auxin, cytokinin, JA, and SA. Auxin involvement is further supported by the presence of auxin-related genes in SA187, as discussed in Section A. Cytokinins, which often act in coordination with auxins, were also reprogrammed, while JA and SA are known to intersect with nutrient signaling and influence root architecture and microbial interactions (Vacheron *et al*., 2013; Guan, 2017). These pathways could likely participate and even engage in hormonal crosstalk and may interact with ethylene-dependent and -independent regulation of NRT2 transporters to fine-tune N accumulation. Future studies using hormone mutants and signaling reporters will be needed to clarify the contribution of these additional hormonal networks to SA187-mediated growth promotion for a better understanding of this beneficial interaction.

Taken together, presented findings collectively support a model in which SA187 promotes nitrate accumulation and N-allocation to the shoot through coordinated regulation of HATS expression via both ethylene-dependent and ethylene-independent pathways. *NRT2.5* functions downstream of ethylene signaling in both shoots and roots, while *NRT2.6* shows a dual regulatory pattern; ethylene-dependent in the shoot, but ethylene-independent in the root (Figure 5). This coordinated spatial and hormonal regulatory control could possibly allow SA187-mediated fine-tuning of nitrate accumulation and N-allocation through multiple entry points in the signaling network, enabling improved plant N-status and internal redistribution under low nitrate conditions. These findings advance our understanding of plant–microbe interactions and offer a framework for leveraging beneficial microbes to enhance NUE and promote sustainable reducing the use of N-based fertilizer agricultural practices. Future studies should extend this framework by dissecting the mechanisms further in-depth investigating hormonal crosstalk, N fluxes, functional validation of role of other HATS, and eventually testing translational applications in crops under field conditions as a critical next step.

## Data Availability Statement

The RNA-seq datasets generated in this study have been deposited in the NCBI Gene Expression Omnibus (GEO) under accession number GSE300561. Reviewer access is available using the token sjihseoillclpwn. The data will be made publicly accessible upon publication.

In addition, the data are available in the CATdb database under the project name *2023_15_QS_SA187* (https://catdb.ips2.universite-paris-saclay.fr/), currently under private access and to be released post-publication.

## Acknowledgments and funding statement

Amina Ilyas was supported by a PhD scholarship from the French Ministry of Higher Education and Research (Ministère de l’Enseignement supérieur et de la Recherche). This work was supported by funding from the Indo-French Centre for the Promotion of Advanced Research (IFCPAR/CEFIPRA) under project 68T06-1 and Saclay Plant Sciences (ANR-17-EUR-0007) Transcriptomics analyses were performed in collaboration with the POPS platform (IPS2, University Paris-Saclay). Seeds of the *nrt2.5*, *nrt2.6*, and *nrt2.5×nrt2.6* mutant lines were kindly provided by Dr. Guilhem Debrosses (Université de Montpellier). The authors would like to thank all colleagues and collaborators who contributed to discussions and technical support throughout the course of this study.

## Author Contributions

A.I., M.H. and A.d.Z. designed experiments, analyzed data and interpreted results. A.I. performed all experiments except for the RNA-seq. A.I. and A.d.Z. participated in supervision. C.M. assisted with plant phenotyping, conducted elemental analysis and participated in the interpretation of the results. J.C. and C.P.-L. conducted RNA-seq and carried out initial analyses respectively. B.D. participated in bacterial quantification assay, *in silico* analysis and phenotyping. A.d.Z. supervised the project, and secured funding. A.I wrote the manuscript and prepared the figures, A.d.Z. and M.H. participated in manuscript revisions.

## Author Approvals

All authors have seen and approved the final version of the manuscript. The work described has not been published previously and is not accepted for publication elsewhere.

## List of Figures

**Figure 1.** The beneficial effect of SA187 on *Arabidopsis thaliana* plant growth parameters under high and low nitrate

**Figure 2.** The effect of SA187 on *Arabidopsis thaliana* shoot and root C and N composition under high and low nitrate

**Figure 3.** Global transcriptomic responses to SA187 in *Arabidopsis thaliana* shoots and roots under high and low nitrate

**Figure 4.** Effect of SA187 inoculation on *Arabidopsis thaliana* wild-type (Col-0) and mutant seedlings grown under low nitrate

**Figure 5.** Functional and in silico analyses of ethylene and nitrate transporter components involved in SA187-mediated growth promotion

**Figure 6.** Simplified model of the regulatory influence of SA187 on the expression of *Arabidopsis thaliana* high-affinity nitrate transporter genes in shoots and roots

## Supplemental information

**Supplemental Figure S1.** Differential gene expression in *Arabidopsis thaliana* under high or low nitrate and with or without SA187 inoculation

**Supplemental Figure S2.** SA187 colonization in Arabidopsis shoots and roots under contrasting nitrate conditions.

**Supplemental Figure S3.** SA187 colonization across Arabidopsis genotypes impaired in ethylene signaling or nitrate transport compared with wild type.

**Supplemental Figure S4.** Effect of SA187 inoculation on Arabidopsis thaliana wild-type (Col-0) and mutant seedlings grown under high nitrate (20mM).

**Supplemental Table S1.** qRT-PCR Primer sets used in this study, related to Figure. 1 and Figure. 4

**Supplemental Table S2**. Differentially expressed genes (DEGs) in shoot tissues of *Arabidopsis thaliana* under 1 mM and 20 mM nitrate with or without SA187 inoculation and GO-term enrichment analyses, related to Figure 3

**Supplemental Table S3.** Differentially expressed genes (DEGs) in root tissues of *Arabidopsis thaliana* under 1 mM and 20 mM nitrate with or without SA187 inoculation and GO-term enrichment analyses, related to Figure 3

**Supplemental File W1.** Analysis of *NRT2.5* and *NRT2.6* promoter using computer simulations. The potential binding sites for EIN3, EIL2 and EIL3 in the promoter of the *NRT2.5* gene (AT1G12940) are identified. The transcriptions binding sites for EIN3, EIL2, and EIL3 are highlighted in the colors red, green, and blue, respectively. The potential binding sites for EIN3, EIL2 and EIL3 in the promoter of the *NRT2.6* gene (AT3G45060) are identified. The transcriptions binding sites for EIN3, EIL2, and EIL3 are highlighted in the colors red, green, and blue, respectively. The analysis was carried out using plantpan 4.0 (Chow et al., 2024) with the standard parameters. The promoters were analyzed 2000bp upstream of the Transcriptional Starting Site (TSS).

