## Supplemental figures for "*Enterobacter* sp. SA187-induced coordinated regulation of high-affinity nitrate transporters and ethylene signaling enhances nitrogen content and plant growth under low nitrate"

### Slide 1
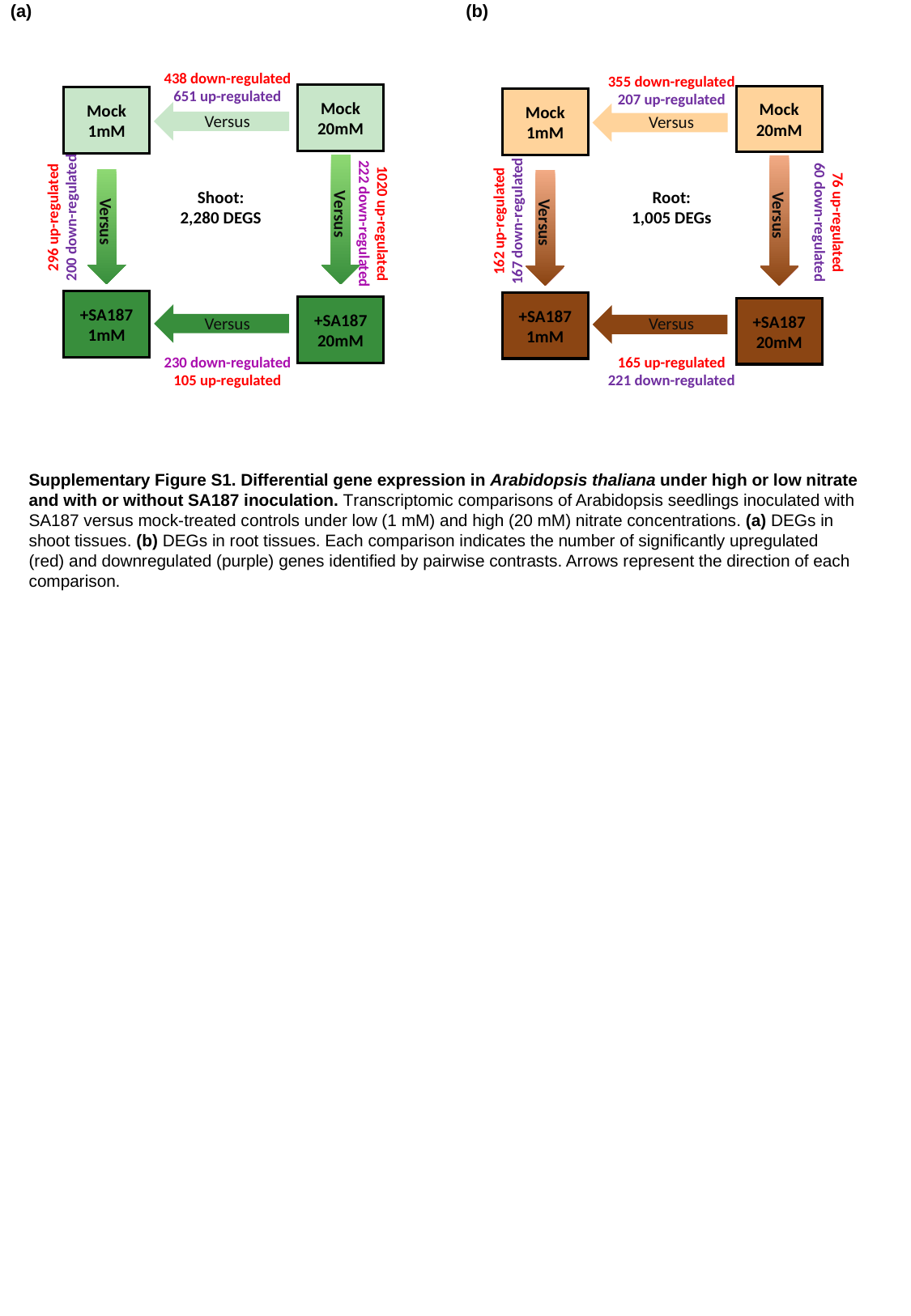

(b)
(a)
438 down-regulated
651 up-regulated
355 down-regulated
207 up-regulated
Mock
20mM
Mock
1mM
Versus
Versus
+SA187
1mM
+SA187
20mM
Mock
20mM
Mock
1mM
Versus
Versus
+SA187
1mM
+SA187
20mM
Versus
Versus
Shoot:
2,280 DEGS
Root:
1,005 DEGs
296 up-regulated
200 down-regulated
162 up-regulated
167 down-regulated
76 up-regulated
60 down-regulated
1020 up-regulated
222 down-regulated
Versus
Versus
230 down-regulated
105 up-regulated
165 up-regulated
221 down-regulated
Supplementary Figure S1. Differential gene expression in Arabidopsis thaliana under high or low nitrate and with or without SA187 inoculation. Transcriptomic comparisons of Arabidopsis seedlings inoculated with SA187 versus mock-treated controls under low (1 mM) and high (20 mM) nitrate concentrations. (a) DEGs in shoot tissues. (b) DEGs in root tissues. Each comparison indicates the number of significantly upregulated (red) and downregulated (purple) genes identified by pairwise contrasts. Arrows represent the direction of each comparison.

### Slide 2
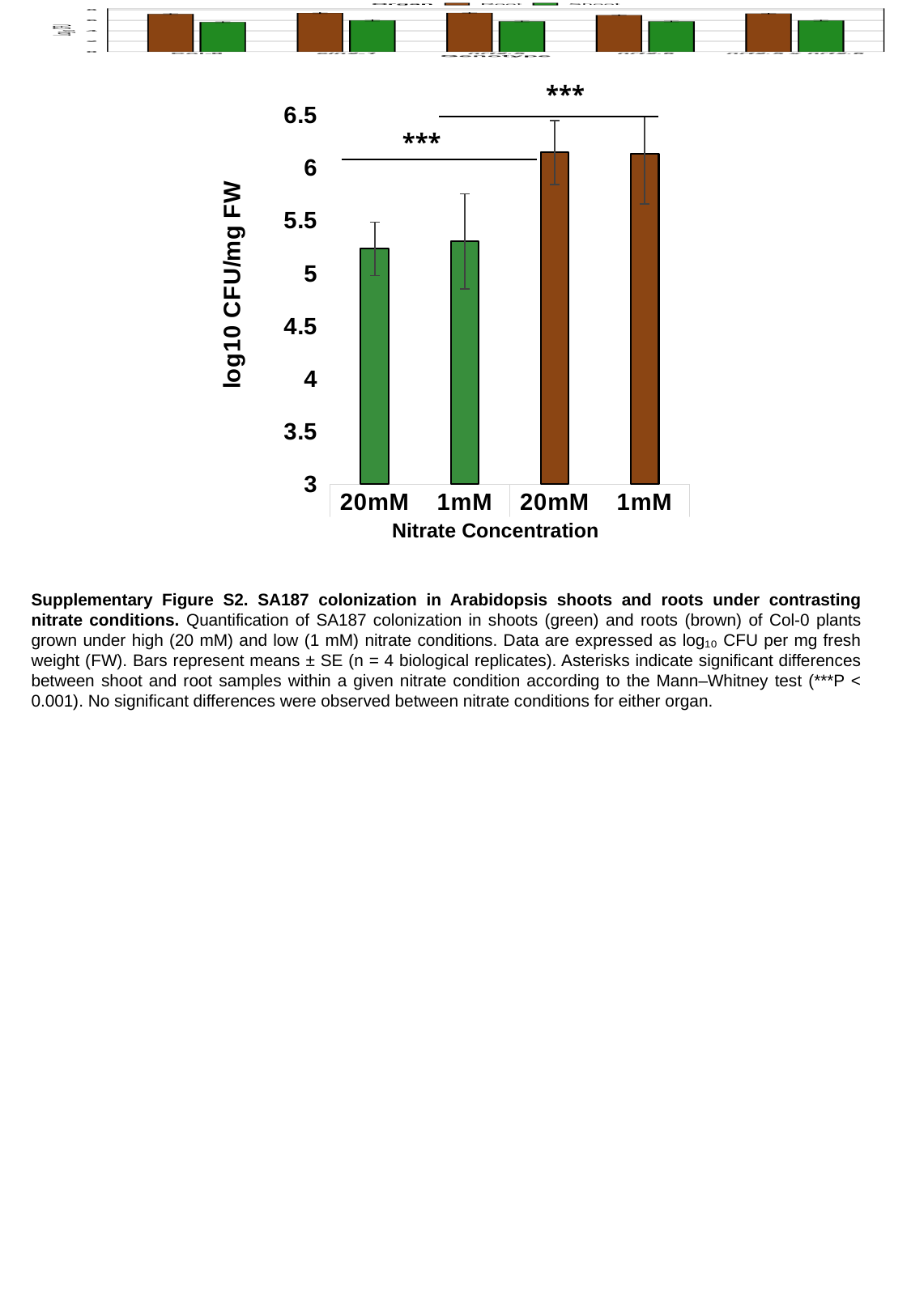

***
#### Chart
| Category | |
|---|---|
| 20mM | 5.234828907043729 |
| 1mM | 5.30452370185004 |
| 20mM | 6.147769969769307 |
| 1mM | 6.134719549525407 |***
Nitrate Concentration
Supplementary Figure S2. SA187 colonization in Arabidopsis shoots and roots under contrasting nitrate conditions. Quantification of SA187 colonization in shoots (green) and roots (brown) of Col-0 plants grown under high (20 mM) and low (1 mM) nitrate conditions. Data are expressed as log₁₀ CFU per mg fresh weight (FW). Bars represent means ± SE (n = 4 biological replicates). Asterisks indicate significant differences between shoot and root samples within a given nitrate condition according to the Mann–Whitney test (***P < 0.001). No significant differences were observed between nitrate conditions for either organ.

### Slide 3
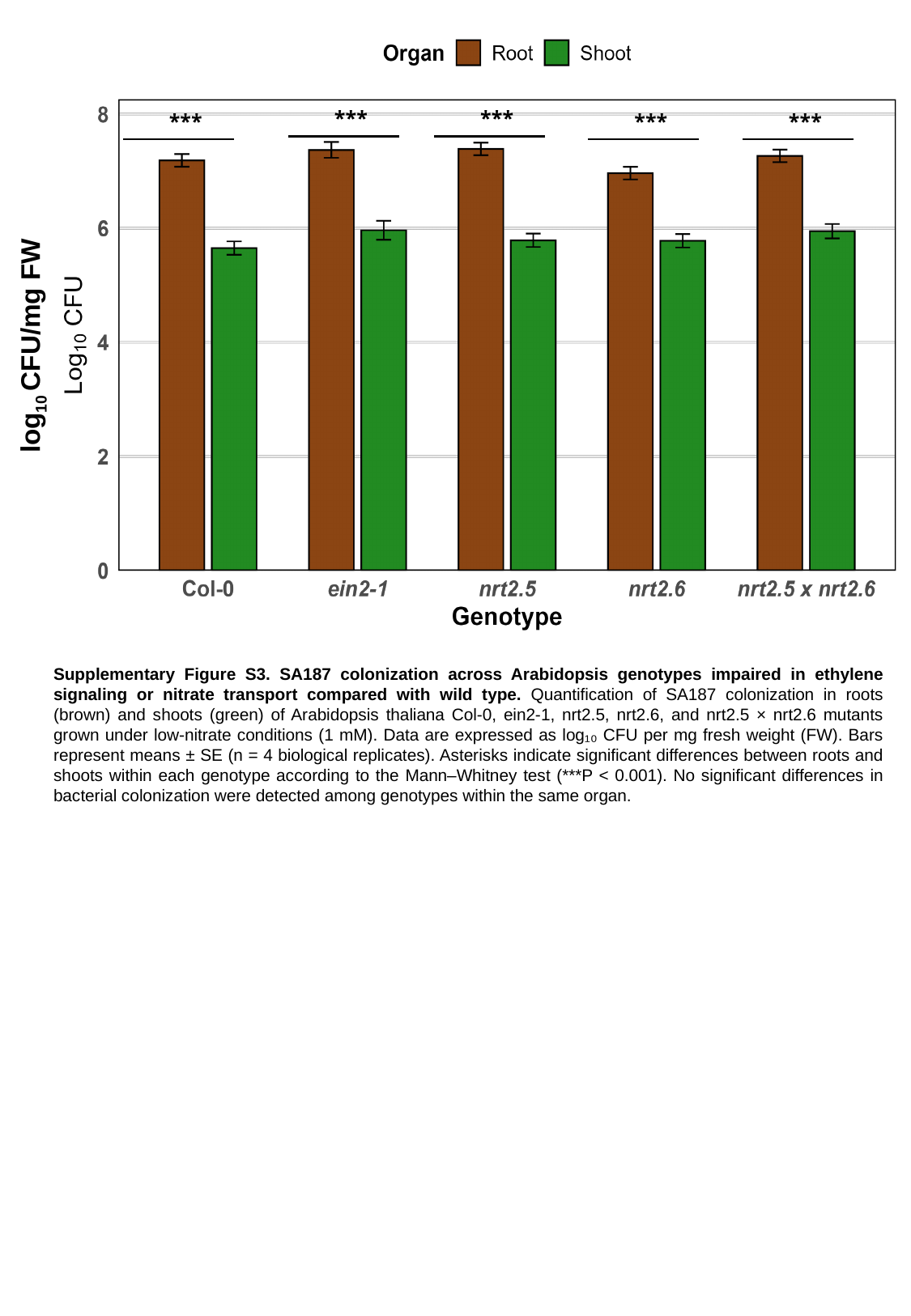

***
***
***
***
***
log_10_ CFU/mg FW
Supplementary Figure S3. SA187 colonization across Arabidopsis genotypes impaired in ethylene signaling or nitrate transport compared with wild type. Quantification of SA187 colonization in roots (brown) and shoots (green) of Arabidopsis thaliana Col-0, ein2-1, nrt2.5, nrt2.6, and nrt2.5 × nrt2.6 mutants grown under low-nitrate conditions (1 mM). Data are expressed as log₁₀ CFU per mg fresh weight (FW). Bars represent means ± SE (n = 4 biological replicates). Asterisks indicate significant differences between roots and shoots within each genotype according to the Mann–Whitney test (***P < 0.001). No significant differences in bacterial colonization were detected among genotypes within the same organ.

### Slide 4
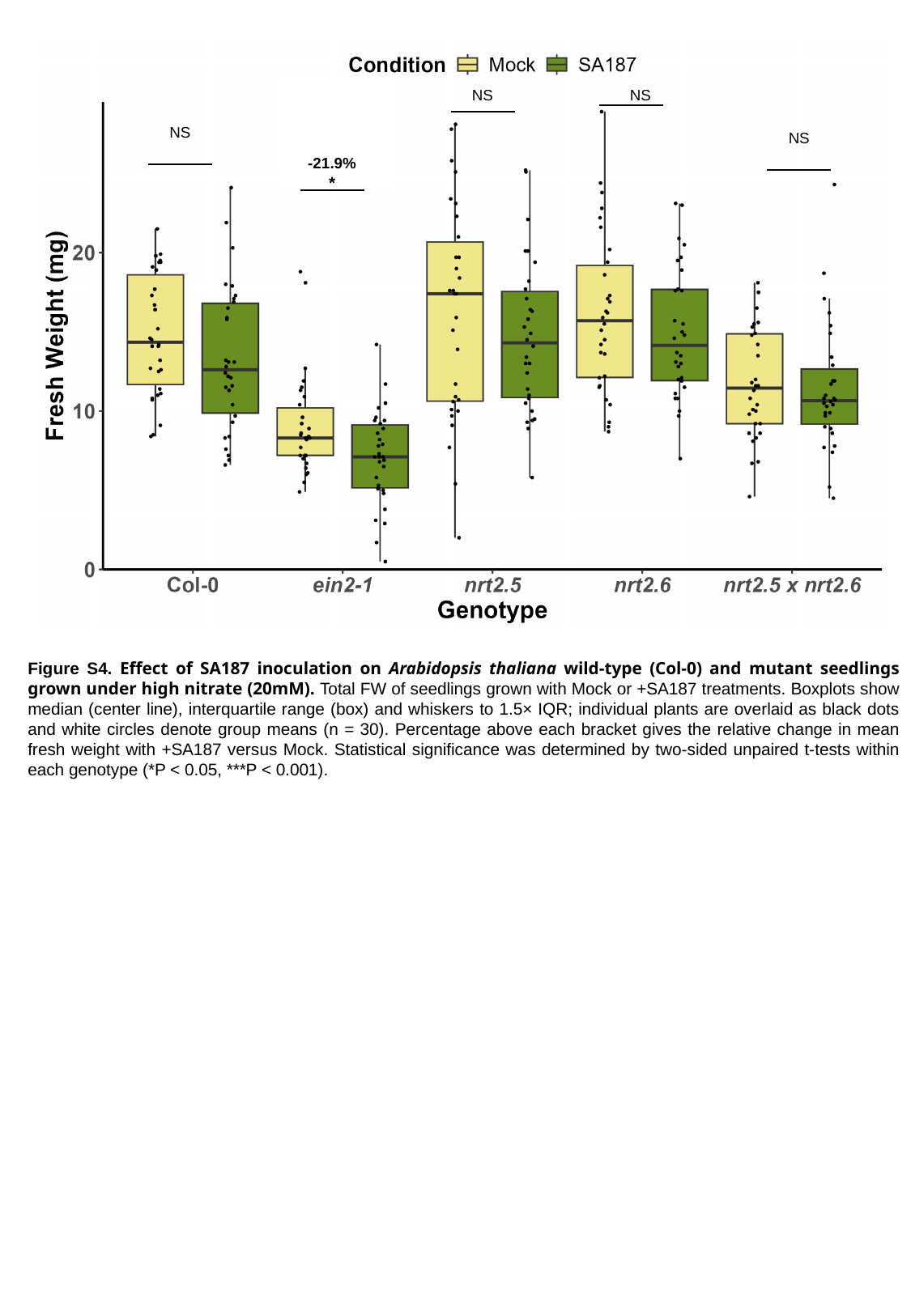

NS
NS
NS
NS
-21.9%
*
Figure S4. Effect of SA187 inoculation on Arabidopsis thaliana wild-type (Col-0) and mutant seedlings grown under high nitrate (20mM). Total FW of seedlings grown with Mock or +SA187 treatments. Boxplots show median (center line), interquartile range (box) and whiskers to 1.5× IQR; individual plants are overlaid as black dots and white circles denote group means (n = 30). Percentage above each bracket gives the relative change in mean fresh weight with +SA187 versus Mock. Statistical significance was determined by two-sided unpaired t-tests within each genotype (*P < 0.05, ***P < 0.001).
