## Supplementary material for "*Enterobacter* sp. SA187-induced coordinated regulation of high-affinity nitrate transporters and ethylene signaling enhances nitrogen content and plant growth under low nitrate": W1

Analysis of *NRT2.5* promoter using computer simulations. The potential binding sites for EIN3, EIL2 and EIL3 in the promoter of the *NRT2.5* gene (AT1G12940) are identified. The transcriptions binding sites for EIN3, EIL2, and EIL3 are highlighted in the colors red, green, and blue, respectively. The analysis was carried out using plantpan 4.0 (Chow et al., 2024) with the standard parameters. The promoters were analyzed 2000bp upstream of the Transcriptional Starting Site (TSS).

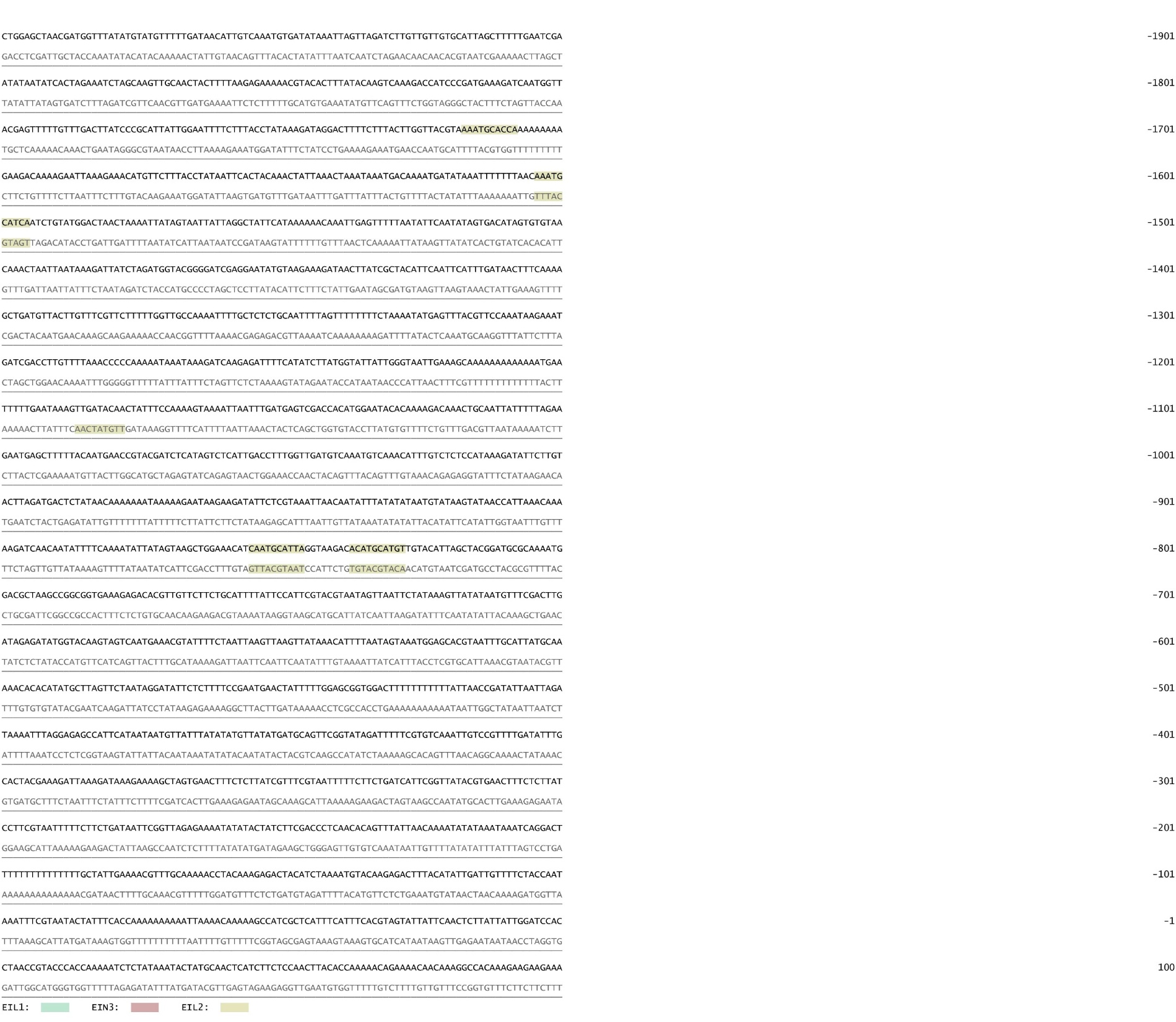

Analysis of *NRT2.6* promoter using computer simulations. The potential binding sites for EIN3, EIL2 and EIL3 in the promoter of the *NRT2.6* gene (AT3G45060) are identified. The transcriptions binding sites for EIN3, EIL2, and EIL3 are highlighted in the colors red, green, and blue, respectively. The analysis was carried out using plantpan 4.0 (Chow et al., 2024) with the standard parameters. The promoters were analyzed 2000bp upstream of the Transcriptional Starting Site (TSS).

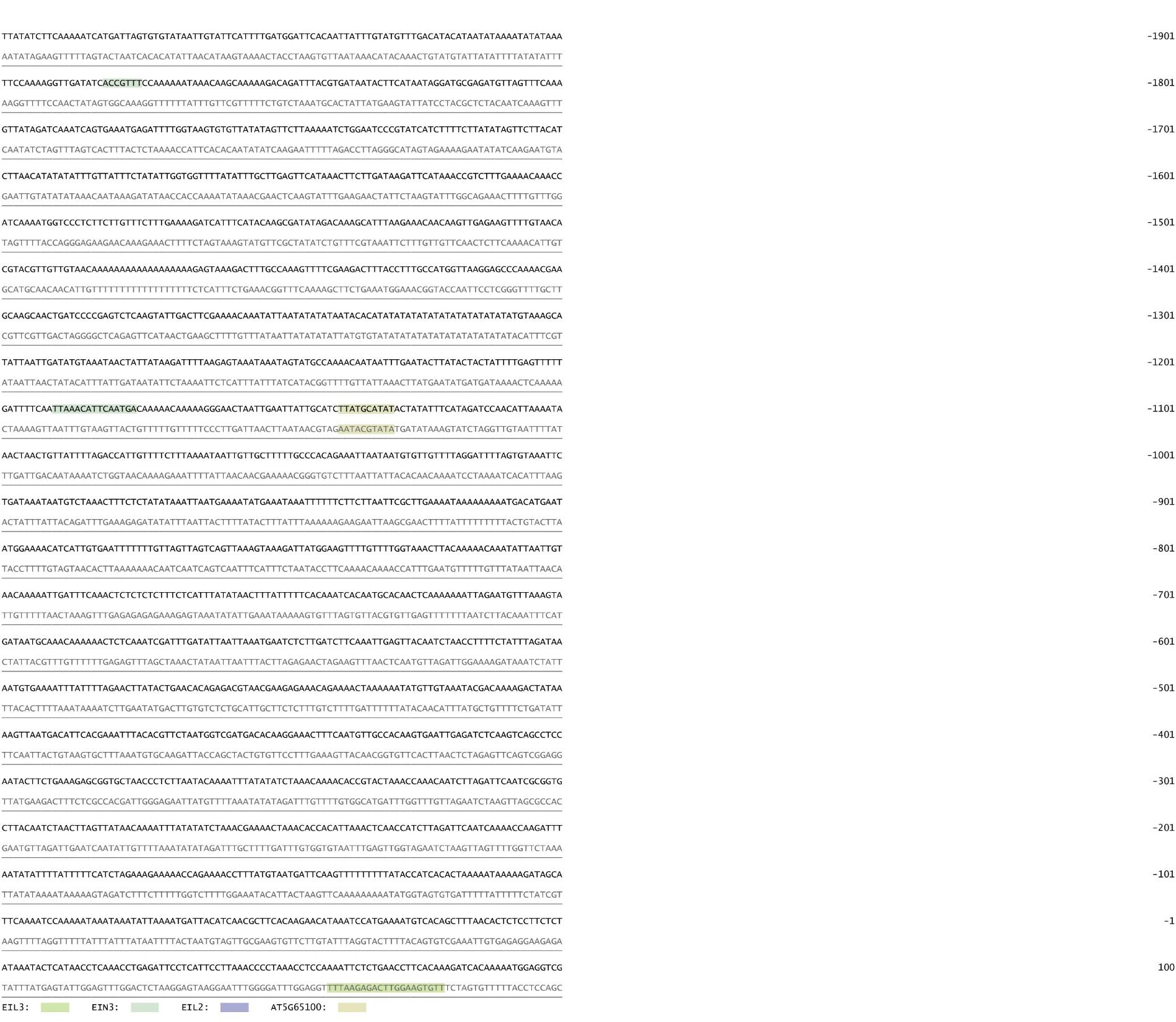

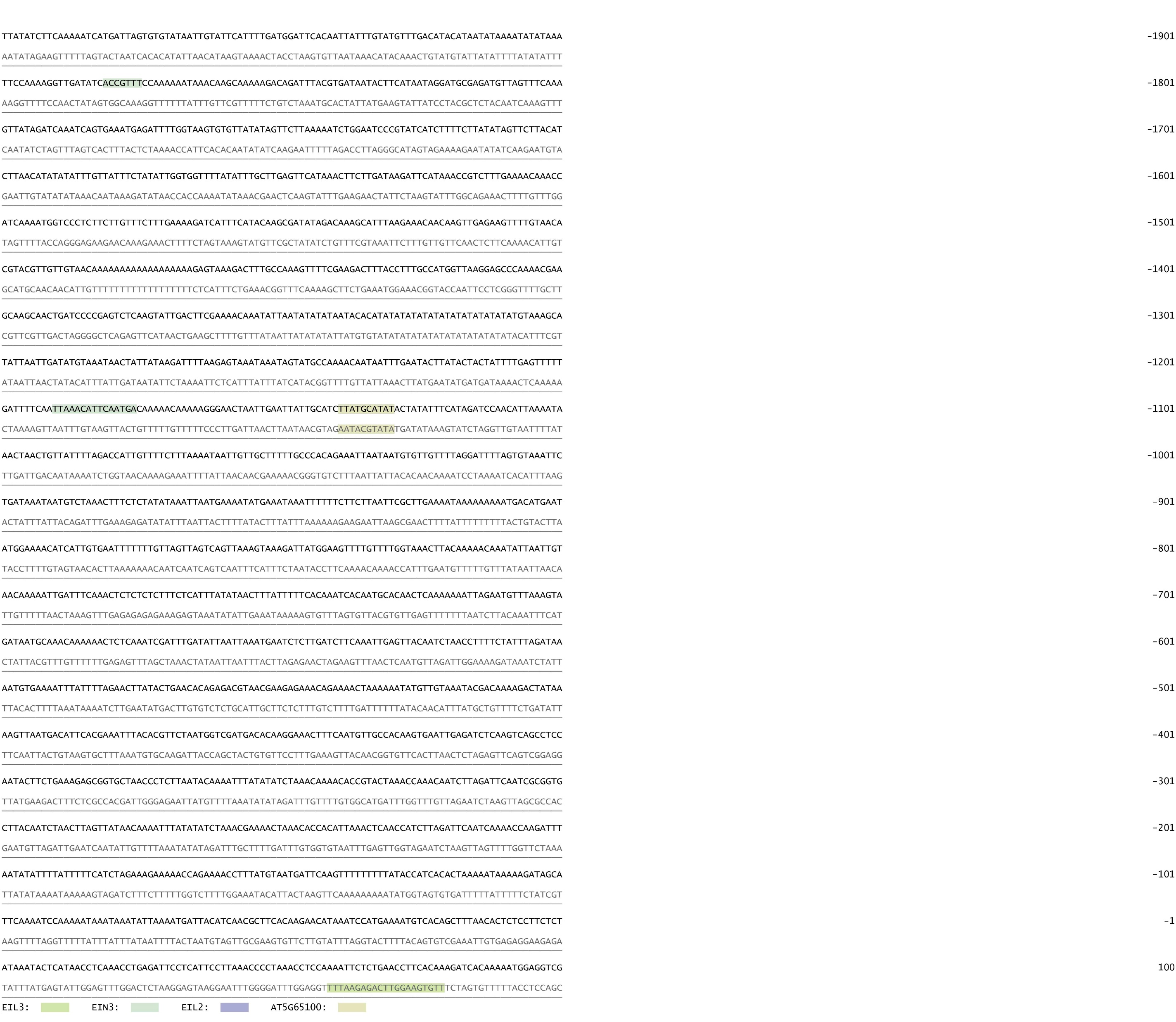
